# PETase Kubu enables near-complete enzymatic depolymerization of commercial PLA/PBAT blend mulch film

**DOI:** 10.64898/2026.06.02.729468

**Authors:** Hong Rae Kim, Hyunjin Kim, Soyeong Jeong, Ju Hee Hwang, Dongsik Kim, Yeon Woo Hong, Dong-Eun Suh, Sukkyoo Lee, Sangmin Lee, Jang-Hee Cho, Jisang Yu, Juntaek Oh

## Abstract

Agricultural mulch films improve crop productivity, but post-use recovery and recycling remain difficult because the films are thin, fragmented, dispersed across fields, and contaminated. Commercial PLA/PBAT blends are increasingly used as biodegradable mulch film materials, yet these films exhibit slow or incomplete degradation under environmental conditions. Here, we show that Kubu, a thermostable PETase from *Kutzneria buriramensis*, rapidly depolymerizes commercial PLA/PBAT mulch film without pretreatment at 60 °C, achieving 95% mass loss within 96 h and releasing terephthalate, adipate, and lactate, detected by LC-MS/MS and HPLC, as the major monomeric products of both PBAT and PLA components. GPC and SEM revealed extensive degradation at the polymer-water interface. DiffDock docking against Kubu, IsPETase, and TfCut indicates that the canonical W/F(Y) cleft of the PETase/Cutinase fold accommodates aliphatic and aromatic ester bonds with comparable geometry, suggesting that substrate promiscuity is an inherent property of the cleft architecture. Kubu’s distinctive contribution combines this permissive cleft with catalytic activity sufficient for near-complete blend depolymerization within 24 h. Integration with a socioeconomic analysis shows that complete enzymatic depolymerization could avoid social costs up to $4,087 per ton of mulch film. These findings establish a single-enzyme approach to end-of-life management of heterogeneous polyester blends.

## Introduction

Climate change threatens global food security, mainly by lowering crop yields through greater exposures to extreme weather^1-3^. Soil management using plastic film mulching is considered as a potential climate change adaptation pathway as it helps retaining soil moisture and reducing pests and diseases^4,5^ (Fig. 1). With its yield-enhancing feature and greater water use efficiency combined with relative low material costs, plastic mulching has been widely adopted globally. While precise relevant statistics are lacking, the global estimate of mulch film usage is about 2.5 million tonnes per year^6^.

**Figure 1.**
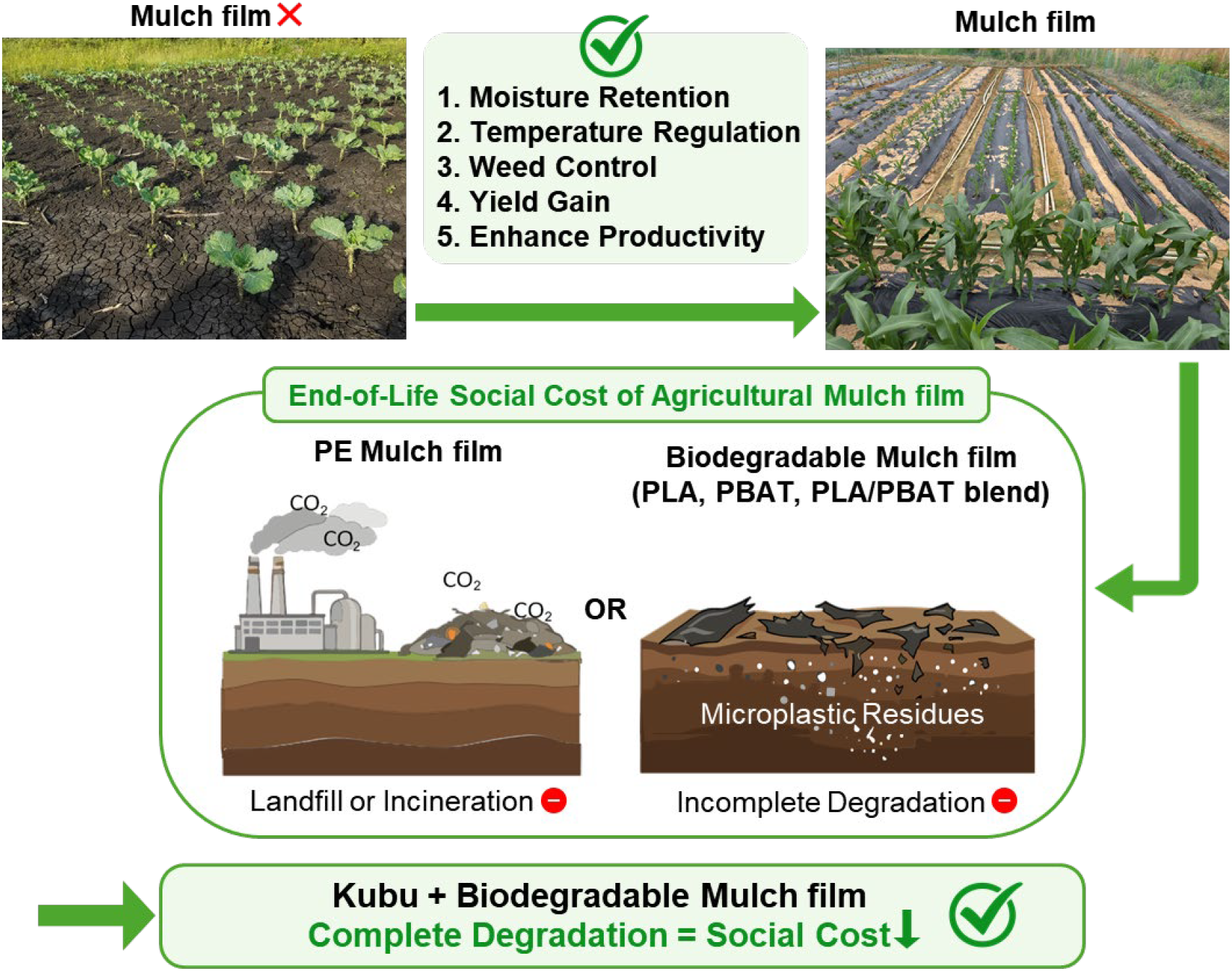
End-of-life social cost of agricultural mulch film and Kubu as a potential enzymatic solution for PLA/PBAT blend mulch film degradation. Schematic illustration of the value of mulching and the comparison of end-of-life pathways for agricultural mulch films and their associated social costs. Conventional polyethylene (PE) mulch films undergo landfill or incineration, generating greenhouse gas and toxic emissions with social costs. Biodegradable mulch films undergo incomplete in-soil degradation, releasing microplastic residues with social costs. Enzymatic depolymerization of PLA/PBAT blend mulch film by Kubu, demonstrated in this study, achieves near-complete degradation and recovers monomeric building blocks for chemical recycling.

Polyethylene, which comes from non-renewable and fossil-fuel based sources, dominates the commercial usage of mulching raising concerns on “White Pollution”, i.e., generating greenhouse gas emissions and microplastic pollutions throughout the life cycle of the product. Biodegradable films, thus, have been considered as sustainable alternatives. Yet, incomplete degradation and microplastic residues can counteract reversing the promised sustainability (Fig. 1).

Among biodegradable polyesters, PLA and PBAT are frequently blended in commercial mulch films because their mechanical properties are complementary. PLA is a biomass derived polyester with high stiffness but limited toughness, and PBAT is a ductile aliphatic-aromatic copolyester whose tensile strength is insufficient for film use alone^7-9^. Blending the two yields flexible films that meet the mechanical demands of agricultural deployment^10^. However, blending also creates a chemically heterogeneous solid substrate. The film contains three classes of ester linkage. PBAT contains aromatic terephthalate-butylene and aliphatic adipate-butylene esters, and PLA contains lactate esters. Although both polymers are used as biodegradable polyester materials, commercial PLA/PBAT blend films often show incomplete degradation under agricultural field conditions. Field studies have reported fragmentation of biodegradable mulch films rather than complete depolymerization, with residual microplastic fragments persisting in soil after repeated seasonal exposure^11-13^. For example, a recent field study has shown that PLA and PBAT-based biodegradable mulch films only degraded by 46.2% and 88.1% and produced massive microplastics^14^. Microbial consortia may in principle address this chemical complexity through division of labor among complementary enzymes, but such processes are difficult to define, optimize, and translate into controlled depolymerization system^15^. These limitations highlight the unmet need for defined biocatalysts, ideally a minimal enzyme set or a single enzyme, that can depolymerize intact PLA/PBAT blend films by attacking three distinct ester linkages within the same solid substrate.

Most of the progress in enzymatic polyester depolymerization has been focused on chemically homogeneous substrates. The discovery of *Ideonella sakaiensis* PETase established that bacterial esterases can hydrolyze PET at moderate temperatures, and rational engineering of the leaf-branch compost cutinase yielded a variant that achieves 90% PET depolymerization within 10 h, advancing towards industrial scale^16,17^. Engineered cutinases from *Thermobifida fusca* convert PBAT films into terephthalic acid (TPA) rich soluble products within 48 h^18^. PLA has been hydrolyzed by proteinase K since the foundational report of Williams in 1981, and microbial cutinases and esterases that act on PLA have also been characterized^19-21^. However, each of these successes addresses a chemically homogeneous substrate. Research on a single enzyme that depolymerizes a heterogeneous PLA/PBAT blend has been limited.

Kubu, a thermostable PETase from *Kutzneria buriramensis* recently identified through global landscape profiling of natural PETase sequences, offers a mechanistically interesting entry point into this gap^22^. Although Kubu was discovered and characterized as a PET hydrolase, its broad and open active site architecture suggests potential compatibility with chemically distinct polyester substrates, a possibility that has not yet been tested. PETase family enzymes have historically been described as aromatic polyesterases incompatible with aliphatic substrates, and whether Kubu also inherits this canonical specificity is unknown^23^.

In this study, we show that Kubu, previously known as PETase, rapidly depolymerizes a commercial PLA/PBAT mulch film without physical or chemical pretreatment under aqueous conditions at 60 °C. The reaction proceeds to near-complete depolymerization within 24 h and releases terephthalate, adipate, and lactate, the three monomeric building blocks of the blend, in quantities consistent with full conversion of both polymer components. We combine film mass loss assays, response surface optimization, apparent Michaelis-Menten kinetics, LC-MS/MS and HPLC product analysis, gel permeation chromatography, scanning electron microscopy, and plastic oligomer docking using DiffDock against the wild-type Kubu structure to define the catalytic performance and structural basis of Kubu mediated PLA/PBAT depolymerization compared to that of PET. We then quantify the social value of complete enzymatic depolymerization from avoided social cost from using PE mulching (greenhouse gas emissions) or from incomplete avoided microplastic burden. Together, these results establish Kubu as a single enzyme to process commercial PLA/PBAT mulch film to recoverable monomers, and identify the active site feature that allows a PETase scaffold to overcome the chemical heterogeneity of the blend.

## Results

### Kubu degrades commercial PLA/PBAT mulch film without any pretreatment

To test whether Kubu can depolymerize a chemically heterogeneous PLA/PBAT blend in its commercial form, we incubated commercially available PLA/PBAT (50:50 w/w) mulch film with purified wild-type Kubu in aqueous buffer (see SI method section for detail). Throughout this study we used wild-type Kubu rather than the engineered variant Kubu-P^M12^ reported in the same work^22^. Kubu-P^M12^ was selected for ethylene glycol tolerance under high temperature glycolysis conditions, a chemistry distinct from the aqueous hydrolysis of PLA/PBAT examined here, and the wild-type enzyme represents the intrinsic substrate envelope of the Kubu scaffold. The film was used directly in its original form without grinding, melt processing, or chemical pretreatment. The lipase PFL1 from *Pelosinus fermentans*, previously reported to act on polyethylene and PBAT, was screened in parallel as a comparator^24^. PFL1 showed substantially lower activity than Kubu under all conditions tested (Fig. S1-S3, see SI for detail). Therefore, we focused on wild-type Kubu in this study.

In our assay condition, wild-type Kubu was sufficient to depolymerize the intact commercial PLA/PBAT mulch film, suggesting that the native PETase scaffold can attack the chemically heterogeneous blend substrate (Fig. 2). This reaction was strongly temperature dependent. Mass loss measurements over 96 h showed that degradation efficiency was maximum at 60 °C with comparable activity also observed at 45 °C and 75 °C but minimal activity at 30 °C. This optimum likely reflects a balance between enzyme activity and the temperature-dependent physical state of the solid substrate. Because the glass transition of PLA typically occurs near 60 °C, increased segmental mobility of PLA-rich amorphous regions may improve access of Kubu to ester bonds within the blend film. The reduced activity at 75 °C suggests partial thermal denaturation or reduced activity of enzyme under non-optimal temperature. Thus, the 60 °C optimum likely arises from combined effects of substrate accessibility and enzyme thermostability, rather than from catalytic temperature dependence alone. It is noteworthy that low Kubu concentration of 0.05 and 0.10 mg/ml produced more efficient mass loss, whereas higher concentrations of 0.20 and 0.40 mg/ml reduced activity (Fig. S2). This pattern suggests surface crowding or nonproductive adsorption, phenomena reported for PETase reactions on solid polyester substrates^25^. We therefore used 0.10 mg/mL as the standard Kubu concentration for subsequent assays to provide robust product signals while remaining within the high activity concentration range. Under this condition at 60 °C, the PLA/PBAT mulch film underwent extensive fragmentation, with 86% mass loss within 24 h and 95% mass loss after 96 h (Fig. 2b). The PFL1 and Kubu combination consistently underperformed Kubu alone across all plastic loadings (Fig. S3), further supporting this competitive nature of two enzymes, possibly through competitive binding on the film surface. All subsequent characterization was therefore performed with Kubu alone.

**Figure 2.**
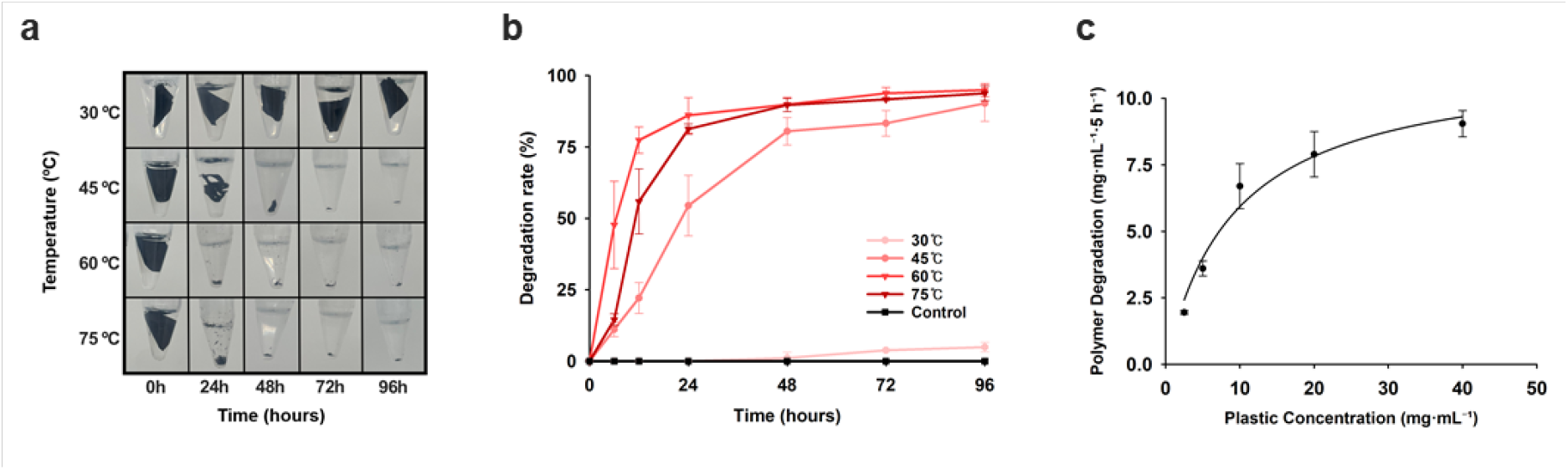
Temperature-dependent degradation of PLA/PBAT blend films by Kubu. (a) Time-course degradation of PLA/PBAT blend films by Kubu at different temperatures. (b) Degradation efficiency of PLA/PBAT blend films as a function of temperature and reaction time for Kubu (n=3). (c) Amount of degraded plastic as a function of substrate concentration.

To measure quantitative activity of Kubu on the solid PLA/PBAT substrate, we analyzed 5 h mass loss at 60 °C using various substrate loading from 2.5 to 40 mg/ml with a Kubu concentration of 0.10 mg/ml (Fig. 2c). As substrate loading increased, the absolute mass of degraded polymer increased toward a plateau. Michaelis-Menten analysis yielded apparent kinetic parameters, Km_app of 9.37 (mg·mL^−1^) and Vmax_app of 11.49 (mg·mL^−1^) per 5h (Table S1). Although these values cannot be interpreted as kinetic values, the apparent Km and Vmax provide a useful benchmark for evaluating future improvements to the enzyme.

### Reaction optimization and identification of monomeric end products

The optimal reaction conditions for Kubu-mediated degradation of the PLA/PBAT blend were determined using response surface methodology (RSM)^26,27^. We applied RSM with three factors: temperature (52.5-67.5 °C), pH (7.0-9.0), and agitation rate (0-200 rpm), with mass loss after 18 h as the response (Table S2). The resulting model exhibited statistical significance with an R^2^ value of 0.941. Among the three key reaction parameters evaluated the optimal conditions were identified as pH 7.4, 60 °C, and 200 rpm with predicted maximum mass loss of 89.5% (Fig. 3). Both pH and temperature exerted a pronounced influence on the degradation rate, whereas rpm showed relatively lower significance within the tested range. To validate the model predictions, additional experiments were conducted under the predicted optimal conditions. The experimentally measured degradation efficiency at pH 7.4 was 86.9%, which was higher than those observed at pH 7.0 and pH 8.0, thereby confirming pH 7.4 as the true optimum and supporting the reliability of the RSM-based model (Fig. 3c and 2d).

**Figure 3.**
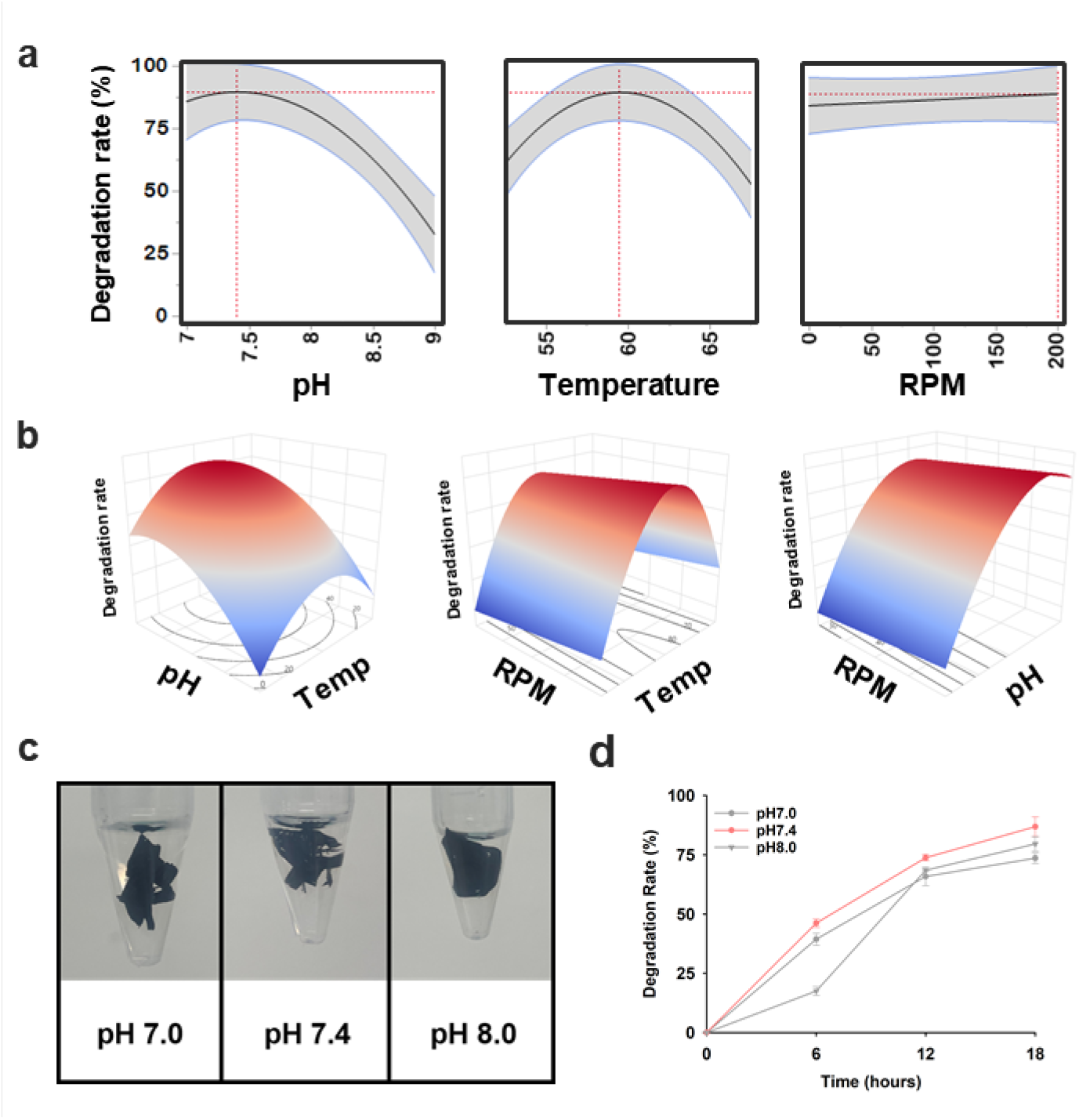
Response surface modeling and optimization of reaction conditions. (a) Predicted effects of key reaction parameters on degradation efficiency. (b) Response surface plot illustrating degradation efficiency. (c) Experimental validation of the predicted optimal conditions. The reactions were performed for 6 hours. (d) Degradation efficiency under different pH conditions (n=3). Statistical significance is indicated as p < 0.05 (*), p < 0.01 (**), and p < 0.001 (***).

Under this optimized condition, we identified the chemical end products of Kubu mediated degradation by using LC-MS/MS. Multiple peaks appeared in the supernatant of Kubu treated reactions (Fig. 4a). The two dominant peaks, labeled (i) and (ii), were identified as TPA and adipic acid. These compounds correspond to the major monomeric units released through the hydrolytic cleavage of ester bonds within the PBAT polymer backbone, as indicated by the red and blue ester linkages in the polymer structure (Fig. 4b)^28,29^. Additional minor peaks were attributed to oligomeric species or intermediates such as butylene terephthalate. Because lactic acid, the monomer of polylactic acid, exhibits high polarity and low molecular weight, its detection by LC-MS was limited^9^. Therefore, its concentration was quantitatively determined using HPLC with a lactic acid standard. Lactic acid concentrations of 257 mg/L and 261 mg/L were detected in the Kubu- and PFL1-treated samples, respectively (Table S3). These results demonstrate that a single enzyme thus converts a heterogeneous PLA/PBAT blend into three distinct monomeric products that span the chemical diversity of the polymer (terephthalate, adipate, and lactic acid). Each is a commodity feedstock currently produced at industrial scale for polymer manufacture, so enzymatic depolymerization recovers chemically reusable building blocks.

**Figure 4.**
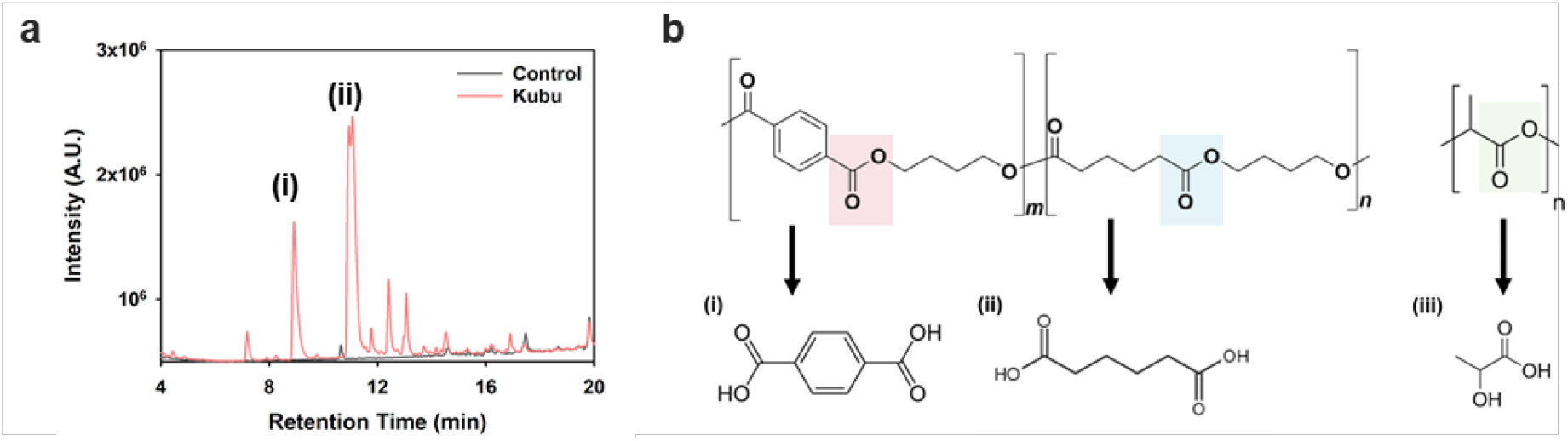
Identification of degradation products. (a) LC–MS/MS chromatograms of degradation products. (b) Chemical structure of the plastic and its corresponding monomeric products.

### Polymer characterization reveals extensive degradation of the polymer surface

To understand how Kubu disassembles the PLA/PBAT film at the polymer level, we tracked average chain length by gel permeation chromatography (GPC) and surface morphology by scanning electron microscopy (SEM). Films were sampled at 0, 4, and 7 h under RSM optimized conditions. Compared with the untreated control, an increasing proportion of low molecular weight polymer species was observed as the reaction time increased. Although an overall shift of the chromatogram toward the low molecular weight region was initially expected, the main peak position remained largely unchanged, while additional signals corresponding to lower molecular weights increased progressively (Figure 4a). This behavior can be attributed to the heterogeneous nature of the reaction, as the water-insoluble plastic undergoes enzymatic hydrolysis predominantly at the surface, whereas polymer chains located in the interior remain largely inaccessible to the enzyme.

Quantitative analysis of molecular weight changes further supported this interpretation. The untreated control exhibited a number average molecular weight (M_n_) of 14,800 and a weight average molecular weight (M_w_) of 71,850. After 4 h of enzymatic treatment which corresponds to the onset of visible film cracking, M_n_ and M_w_ decreased to 6,270 and 64,800, representing reductions of 58% and 10%, respectively (Fig. 5b, S5a). After 7 h, when the film was extensively fragmented, M_n_ and M_w_ further decreased to 2,535 and 46,400, corresponding to reductions of 83% and 35%, respectively. These results clearly demonstrate that Kubu mediated degradation leads to substantial depolymerization of the PLA/PBAT blend^30,31^. Because M_n_ is more sensitive to changes in the low molecular weight fraction, whereas M_w_ reflects alterations in the high molecular weight region, the pronounced decrease in M_n_ indicates a significant accumulation of low molecular weight species during degradation. The polymer dispersity index (PDI) increased continuously with reaction time, indicating that enzymatic depolymerization broadened the molecular weight distribution as a consequence of non-uniform chain scission (Fig. S5b).

**Figure 5.**
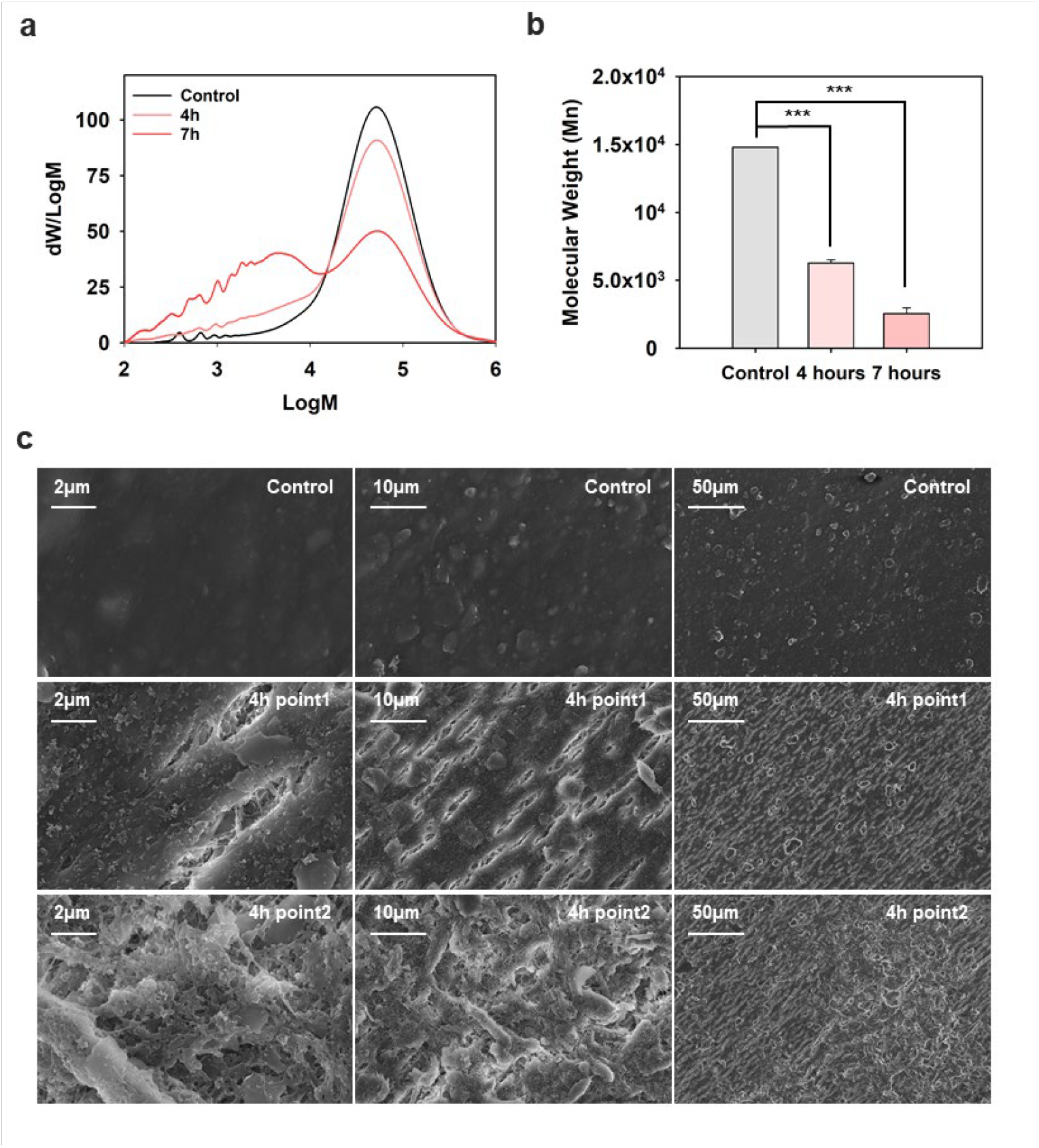
Molecular weight distribution and surface characteristics of enzymatically degraded PLA/PBAT blend films. (a) GPC chromatograms of PLA/PBAT blend films after 4 h and 7 h of Kubu treatment. (b) Changes in number-average molecular weight (M_n_) during enzymatic degradation (n=2). Statistical significance is indicated as p < 0.05 (*), p < 0.01 (**), and p < 0.001 (***). (c) Scanning electron microscopy (SEM) images showing surface erosion of PLA/PBAT blend films after enzymatic degradation.

Surface erosion of the plastic films was further examined by SEM analysis. Only the 4 h-treated samples retained sufficient structural integrity for surface characterization, whereas samples treated for 7 and 10 h were too fragmented for analysis. SEM images revealed that the untreated control exhibited a smooth and uniform surface, while the enzyme-treated samples showed widespread surface cracking (Fig. 5c). At higher magnification, pronounced surface erosion was evident in the Kubu-treated films, providing direct morphological evidence of enzymatically degraded polymers^32^.

### Structural basis for Kubu substrate promiscuity revealed by deep learning-based docking

To understand how Kubu accommodates such chemically diverse ester linkages within a single active site, we constructed a docking ligand library comprising computationally generated PLA and PBAT oligomers representing the ester chemistries present in the PLA blend film, together with BHET (bis(2-hydroxyethyl) terephthalate) as the PET-derived substrate (Fig. 6a, S6 and Table S4). Each ligand was docked against the wild-type Kubu, IsPETase and TfCut structure using DiffDock, a diffusion based generative docking method that has been shown to outperform conventional search based docking for substrates that exceed the typical drug like size range (Fig. 6b)^33^. This feature was important because our goal was to test whether Kubu could accommodate chemically distinct polymer fragments without forcing each substrate into a predefined active site grid. DiffDock samples ligand translation, rotation, and torsional degrees of freedom from random starting configurations and generates multiple plausible poses, making it suitable for comparing flexible polyester oligomers that exceed typical drug like dimensions. We therefore interpreted the docking results not simply by docking score, but by the frequency of reaction compatible poses in which the scissile ester bond was positioned near the catalytic serine residue in an orientation compatible with hydrolysis (see SI methods and Table S5, S6 for details). A higher frequency of such poses was taken as evidence that a given substrate chemistry can be productively accommodated by the Kubu active site. This analysis allowed direct comparison of the native PET substrate with PBAT and PLA ligands present in the commercial blend film (Fig. 6c,d and S7).

**Figure 6.**
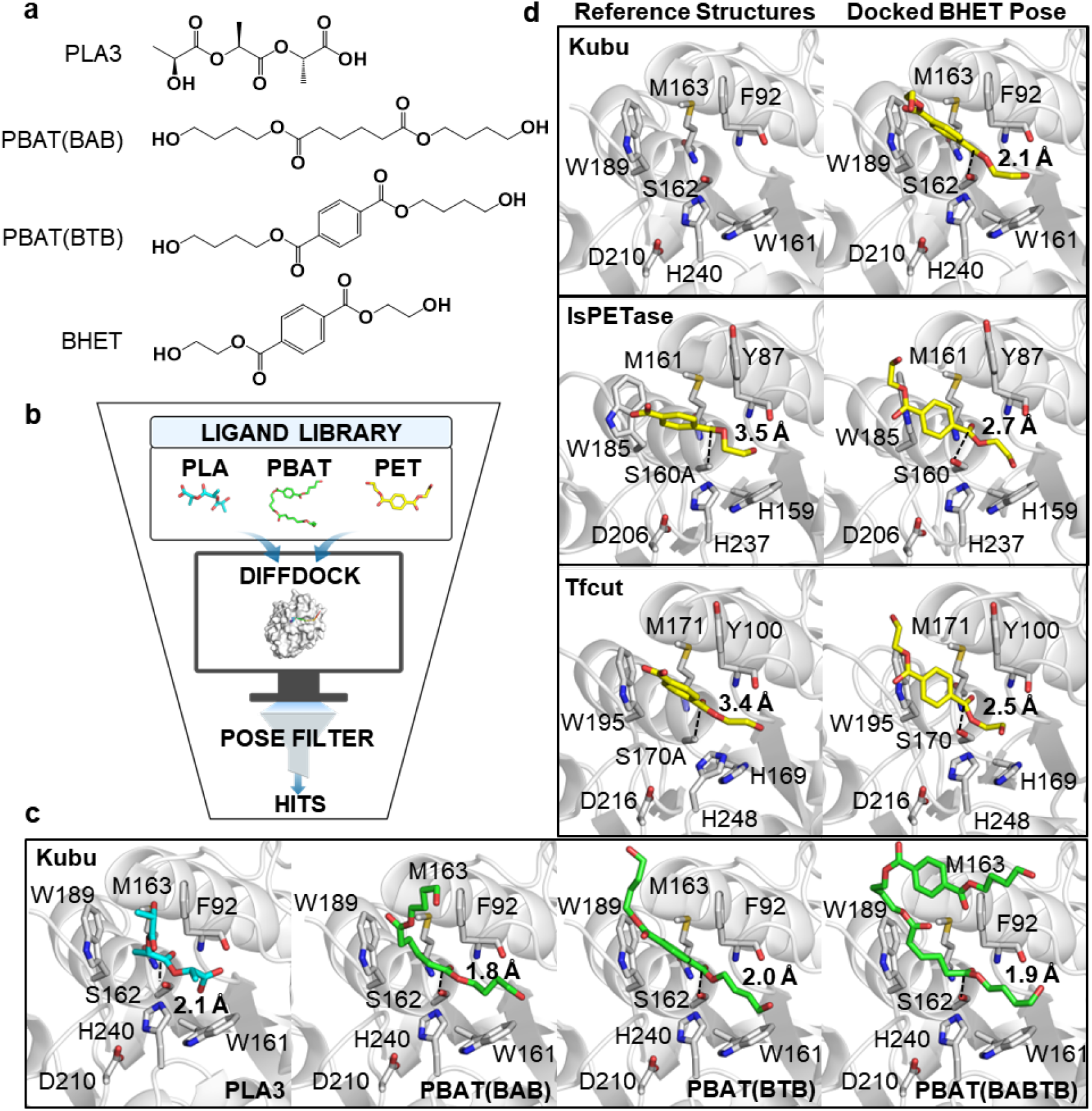
DiffDock docking of a ligand library generated from PLA, PBAT, and PET to Kubu, IsPETase, TfCut. (a) Schematic representation of ligand structures used in docking. (b) Schematic of the DiffDock docking workflow. (c) Representative productive poses of PBAT and PLA oligomers in the Kubu active site. The substrate-binding cleft is formed by F92 and W189. The backbone amides of F92 and M163 form the oxyanion hole. The catalytic triad Ser162-His240-Asp210 is shown. Substrates are shown as cyan, green and yellow for PLA, PBAT and PET, respectively. In all productive poses, the ester-flanking aliphatic carbons of the substrate occupy the cleft between F92 and W189, while the aromatic ring (where present) extends the cleft entrance. (d) Representative productive poses for the BHET docked into IsPETase (PDB 5XJH) and TfCut (PDB 4CG1) active sites^34,35^. Equivalent cleft residues are shown (Y87/W185/M161 in IsPETase; Y100/W195/M171 in TfCut). Apo Kubu (PDB 8YTW) and co-crystal complexes of IsPETase with MHET (PDB 7XTW) and TfCut with MHET (PDB 7XTV) are shown for comparison^18,22^.

In our docking conditions, all three polyester substrate classes (PET, PLA, PBAT) generated productive poses against Kubu at comparable frequencies. PLA dimers through tetramers were each retained as catalytically competent poses, alongside PBAT monomers and dimers spanning both adipate (BAB) and terephthalate (BTB) ester chemistries, as well as the native PET-derived substrate BHET (Fig. 6c,d and Table S7). The retention of productive poses for both purely aliphatic (PLA, PBAT-BAB) and aromatic-containing (PBAT-BTB, BHET) ligands within the same active site indicates that the Kubu cleft accommodates substrate ester linkages of different chemical character without strong geometric discrimination. Inspection of the productive poses shows that Kubu retains the canonical PETase active-site architecture, with the substrate-binding cleft formed by F92 and W189, structurally equivalent to the Y87/W185 cleft of IsPETase and the Y100/W195 cleft of TfCut. The backbone amides of F92 and M163, the residue adjacent to the catalytic Ser162, contribute the oxyanion hole. In productive poses, the ester carbonyl is anchored at this oxyanion hole, with the ester-flanking aliphatic carbons of the substrate extending between F92 and W189; the substrate aromatic ring, where present, is positioned at the cleft entrance rather than sandwiched directly between the cleft residues (Fig. 6c and d).

To test whether this substrate-permissive cleft geometry is unique to Kubu or shared across the PETase/Cutinase fold, we performed parallel DiffDock analysis on IsPETase and TfCut using the same substrate library and pose-filtering criteria^18^. Both enzymes also generated catalytically competent poses for PLA, PBAT, and PET-derived substrates (Table S7, Fig. 6d). The same overall binding geometry, with the substrate aromatic ring extending the cleft entrance and the ester-flanking aliphatic carbons occupying the inner cleft, is also evident in the published IsPETase-MHET and TfCut-MHET co-crystal complexes (Fig. 6c)^18^. The retention of catalytically competent PLA poses by IsPETase, an enzyme reported as inactive on aliphatic polyesters at its 30 °C optimum^23^, suggests that the experimentally reported substrate specificity differences among PETase fold enzymes are not strictly determined by cleft architecture alone. We note that DiffDock generates poses against a fixed receptor structure without explicit binding affinity calculation, and we therefore interpret these docking results as supporting the structural inspection of co-crystal complexes rather than as direct quantitative predictors of catalytic activity.

### Potential value of Kubu in reducing the social cost of mulch film

To deduce the potential value of complete degradation by applying Kubu, we discuss two alternative mulching strategies: traditional PE mulching and using PLA/PBAT blend biodegradable mulching without Kubu and calculate the social costs associated with these two strategies focusing on the end-of-life stage (Fig. 7). Particularly, we estimate the unit social cost that occurs during the disposal stage. For the PE mulching, the cost occurs from the greenhouse gas emission during incineration or landfill. For the biodegradable mulching the cost comes from the microplastic residue stems from incomplete degradation.

**Figure 7.**
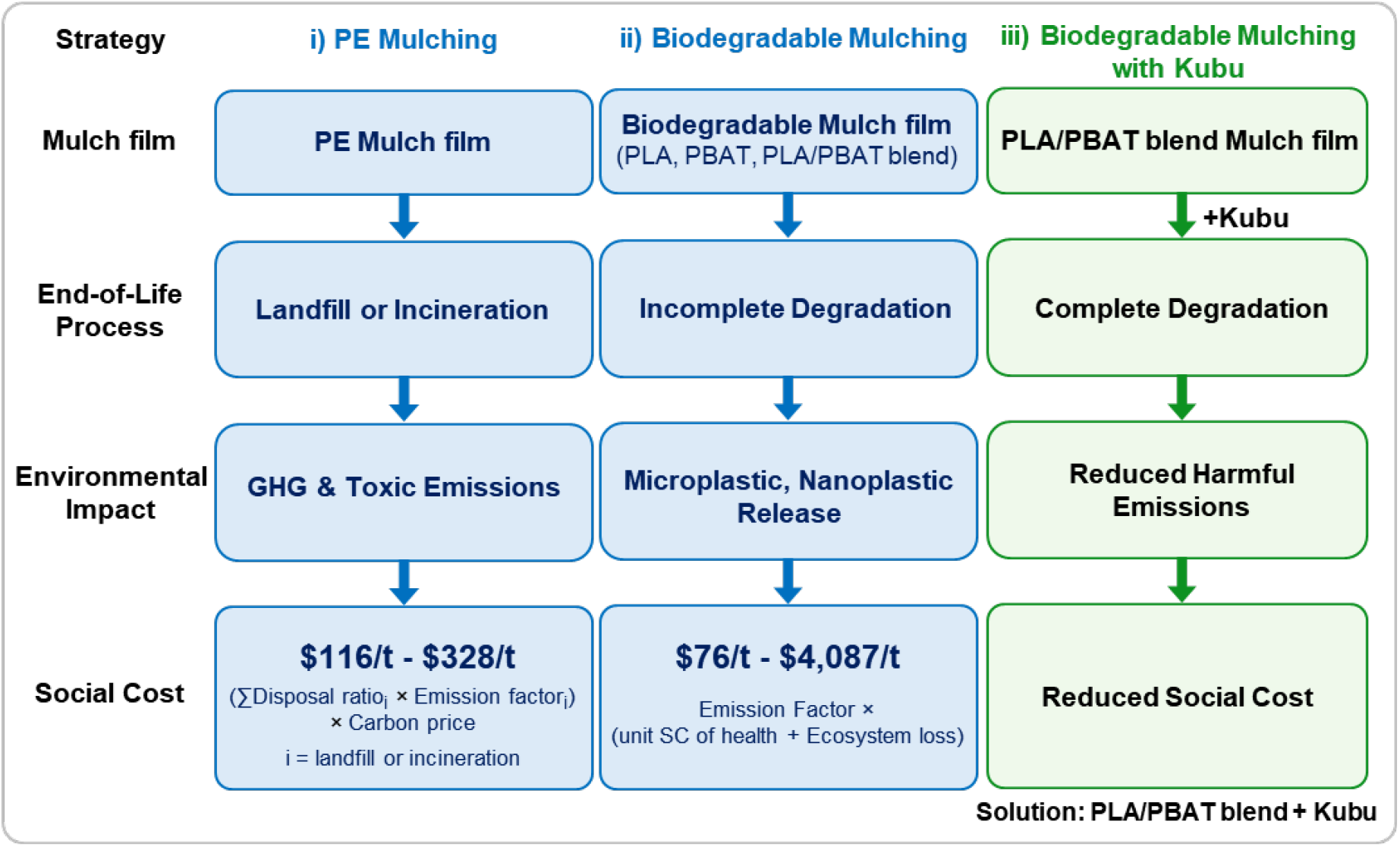
Social cost schematic comparison for End-of-life management of agricultural mulch films. Three alternative strategies of mulching; i) PE mulching, ii) Biodegradable mulching, iii) Biodegradable mulching with Kubu. PE mulching generates GHG and toxic emissions through landfill or incineration, while biodegradable mulching leads to microplastic and nanoplastic release due to incomplete degradation. The strategy of Kubu into PLA/PBAT blend enables complete degradation, resulting in reduced harmful emissions and social cost.

The social cost of using PE mulching can be estimated by using the emission factors for each disposal way reported by Xiong et al. (2023) and Benavides et al. (2020)^36,37^. We apply a weighted average emission factor that reflects the disposal share of each disposal pathway and the emission factor is calculated as 0.96 t*CO*_2_per ton of PE mulch film. Using carbon price scenarios from EPA (2023), based on the Ramsey near-term discount rates, the PE disposal related social cost is ranging from $116/t ∼ $328/t^38^ (Table S8). However, due to data limitations, this estimate does not account for toxic emissions or uncontrolled disposal practices (e.g. open burning), which may generate substantially larger environmental damages. Therefore, the actual social cost of PE disposal could be considerably higher than the estimates presented in this study.

Two potential sources of the social burden from incomplete degradation of biodegradable mulch film are increased *CO*_2_emission from microbial activities and respirations, i.e., priming effect, during degradation, and microplastic release. To our knowledge, such priming effects have not been precisely quantified with a recent study indicating there is no significant priming effects from biodegradable mulch film degradation^39^. More significant burden is from the emission of microplastic. Hence, to estimate social cost from incomplete biodegradation, we use Hao et al. (2024) estimates on the residual microplastic emissions^14^. The study provides estimates to calculate the microplastic emissions per ton of mulch film, which are 0.02t for PLA and 0.11t for PBAT. The social cost of microplastic pollution is from human health damages and ecosystem service losses. Watkins et al. (2022) applied the Disability-Adjusted Life Year (DALY) framework, which measures the loss of healthy life due to premature mortality and morbidity^40^. The health-related social costs are calculated by multiplying the excess disease burden attributable to plastic-related exposure by the monetary value assigned to one DALY. The estimated annual health-related social cost of MNPs is $28.5 billion, consisting of $23.5 billion from direct MNP exposure causing gastrointestinal injury and $5.3 billion from the indirect delivery of other harmful agents by MNPs. The social cost of microplastic per ton is $712.5-$2,850 based on estimated emissions of microplastics to the environment of 10 and 40 million tonnes per year^41^. From an ecosystem perspective, Beaumont et al. (2019) estimated the economic damages of marine plastic pollution based on loss of marine ecosystem services, approximately $500-2,500 billion^42^. Dividing this loss by the estimated global stock of marine plastics in the ocean (75-150 million tonnes) yields an ecosystem-related social cost of approximately $3,300-$33,000 per tonne of marine plastic per year. The unit social cost from microplastic is ranging from $4,013 to $35,850 (Table S9).

In summary, we obtain the estimates for the social cost per ton of mulch film per annum as the followings: PE landfill ($84∼238), PE incineration ($348∼986), PLA microplastic pollution ($76∼681), PBAT microplastic pollution ($457∼4,087). Note that the damages from toxin releases during PE disposal has not been accounted, hence, the social cost of PE usage is underestimated. Also, the more likely pathway to transitioning to biodegradable mulch films is to use the PLA/PBAT blend, which implies that the microplastic releases or the degree of incomplete degradation would be certainly higher than those for PLA films. Finally, we find that even the lower bound estimates are substantial considering the estimated amount of total mulch film usage is around 2.5 million tonnes. Therefore, there is great potential for Kubu to reduce these social costs while the success depends on the cost of applying Kubu.

## Discussion

The ability of a single PETase-family enzyme, Kubu, to release terephthalic acid and adipic acid from the two PBAT ester segments, together with lactic acid from PLA, was unexpected. The canonical view of PETase substrate recognition centers on a hydrophobic active site cleft optimized for aromatic terephthalate, with PET hydrolyzing activity attributed to a “wobbling” tryptophan and adjacent tyrosine forming an aromatic clamp around the terephthalate ring ^16,23,34,43^. The original characterization of IsPETase explicitly reported that the enzyme does not act on aliphatic polyesters, and concluded that PETases are generally aromatic polyesterases^23^. Kubu’s activity on PLA and on the aliphatic adipate-butylene moiety of PBAT therefore extends beyond this canonical specificity and prompts a closer look at the structural basis of polyester hydrolase substrate recognition. A possible structural rationale comes from the docking analysis. PLA and PBAT oligomers were accepted by the Kubu active site at a frequency of catalytically competent poses comparable to that of the native PET substrate (Fig. 6c, d and Table S7), indicating that the cleft does not strongly discriminate among the three ester chemistries present in the blend. In the productive poses, the space between F92 and W189 is occupied by the aliphatic carbons extending from the scissile ester, while the substrate aromatic ring, where present, sits at the cleft entrance rather than stacking directly between the two cleft residues. The same trend is observed in the IsPETase-MHET and TfCut-MHET co-crystal structures, where the terephthalate ring of MHET adopts a tilted orientation that does not align for parallel π-π stacking with either aromatic residue^18^. Together, these structures indicate that productive binding in the W/F(Y) cleft may not strictly require π-stacking between the substrate aromatic ring and the cleft residues.

Under this interpretation, the substrate promiscuity observed for Kubu becomes a predictable property of the cleft architecture rather than a unique feature of this enzyme. PET, PBAT, and PLA all present an ester linkage flanked by short aliphatic chains, and any of these flanking patterns can populate the F92/W189 cleft in catalytically competent geometry. This view is consistent with the substrate breadth of W/F(Y) cleft polyester hydrolases. Cutinases from *H. insolens* and *T. cellulosilytica*, which retain this cleft topology and lack the closed lipase lid, have long been known to act on both aromatic and aliphatic polyesters^44-47^, and engineered *T. fusca* cutinases achieve complete PBAT depolymerization^18^. The W/F(Y) cleft thus appears to confer a broad substrate envelope shared across the PETase/Cutinase fold, with apparent specificity differences among family members reflecting kinetic, thermal factors and other structural diversity rather than cleft breadth only. What distinguishes Kubu from other W/F(Y) cleft polyester hydrolases is therefore not the cleft itself but the conjunction of three properties that together enable near-complete depolymerization of a commercial blend. The open active site shared with cutinases, robust catalytic activity at 60 °C, and catalytic efficiency sufficient to drive the reaction to extensive mass loss within 24 h. This reframes the question of polyester hydrolase substrate specificity from one of cleft architecture to one of cleft compatibility with the polymer being targeted, although extensive mutagenesis and structural study of Kubu, and characterization of other polyester hydrolases that show substrate promiscuity, will be required to establish clear conclusions.

The chemical heterogeneity of commercial PLA/PBAT mulch films has historically been addressed through enzyme combinations or microbial consortia, on the rationale that each polymer chemistry should be matched to its preferred enzyme. Synergistic two enzyme systems have been particularly successful for chemically homogeneous substrates. The canonical PET pathway uses PETase for surface depolymerization and MHETase for hydrolysis of the soluble dimer intermediate, and engineered combinations of these enzymes outperform either alone ^16,48^. Marine and soil microbial consortia handle aromatic-aliphatic copolyesters through division of labor, with different community members expressing complementary hydrolases and downstream catabolic genes^15^. By analogy, a combination of PFL1 (active on aliphatic ester linkages) and Kubu (active on aromatic ester linkages) might be expected to outperform either enzyme alone on a PLA/PBAT blend. However, what we observed was the opposite. Across all substrate loadings tested, the PFL1 and Kubu combination consistently underperformed Kubu alone (Fig. S3). Two factors plausibly contribute to this loss of activity. First, when both enzymes accept the same ester linkages, they compete for a finite number of productive surface attack sites rather than collaborate across chemically distinct sites ^25,49^. Surface saturation is the rate limiting condition for PETase family enzymes acting on solid substrates, and adding a second enzyme that binds the same surface but with lower turnover converts a fraction of productive Kubu substrate complexes into less productive PFL1 substrate complexes^50^. Furthermore, PFL1 as a lipase contains a mobile α-helical lid that occludes its active site in solution^24^, imposing an additional kinetic barrier of lid opening that is absent in the no-lid PETase/Cutinase architecture of Kubu. Second, productive reaction pose of polyester hydrolases requires not only binding but proper orientation of the catalytic triad relative to a surface exposed ester bond. Two enzymes occupying adjacent surface positions may sterically interfere with each other’s productive orientations^51^. Our observations suggest that combining two enzymes is productive when their substrate preferences are non-overlapping and their reaction products feed sequentially, as in PETase to MHETase. When substrate preferences overlap on a heterogeneous solid surface, combination becomes a competitive rather than cooperative arrangement. A single enzyme with sufficiently broad substrate specificity, such as Kubu, bypasses the problem entirely. This reframes the design problem for chemically heterogeneous biodegradable polymers from one of enzyme combination to one of single broad substrate enzyme discovery. Natural sequence space, as demonstrated by the discovery of Kubu, likely contains additional polyester hydrolases whose substrate range have not yet been characterized.

Applying Kubu to achieve fast and complete degradation of biodegradable mulch films can reach substantial reduction in social cost from using mulch films. Our socioeconomic analysis implies that two alternative mulching strategies, traditional PE mulching and biodegradable mulching with incomplete film degradation, have substantial social costs. Per ton of mulch film, the estimates range from $76 to $4,087, which lead to 190 million to 10 billion USD of total annual social cost based on 2.5 million tonnes of mulch film usage. As disposal and recycle through applying Kubu can offset these costs, the key future question is by how much considering the cost of Kubu application and potential GHG emission from film collection.

We have shown that Kubu, a thermostable PETase recently identified^22^, achieves near-complete depolymerization of a commercial PLA/PBAT mulch film within 24 h at 60 °C. The reaction releases the three monomeric building blocks of the blend, terephthalic acid, adipic acid, and lactic acid, in quantities consistent with near-complete depolymerization of the blend. Apparent Michaelis-Menten kinetics, gel permeation chromatography, and scanning electron microscopy together establish that the enzyme generates low molecular weight fragments from the polymer-water interface. DiffDock-based docking against the wild-type Kubu structure indicates that PLA and PBAT oligomers can adopt catalytically competent poses at the active site at a frequency comparable to that of the native PET substrate, providing a candidate structural basis for the observed substrate promiscuity. Socioeconomic analysis suggests complete enzymatic depolymerization avoids both the CO_2_ mitigation foregone from using traditional PE mulching and the microplastic externality that arise from partial in-soil degradation of biodegradable mulching, while recovering monomers as commodity feedstocks for chemical recycling.

The limitation of this work is that Kubu’s catalytic optimum lies at 60 °C. This temperature range is well matched to industrial composting and centralized bioreactor scenarios. However, mulch film application in agriculture spans a wide range of contexts. Recovery infrastructure suitable for centralized enzymatic recycling exists in some agricultural systems but is logistically and economically infeasible in others, particularly in regions where mulch film usage is highest in absolute terms^52,53^. In these contexts, in-soil or near-soil degradation under ambient soil temperatures is the only practical disposal route, and 60 °C optimized enzyme is difficult to be applied. Bridging the gap between laboratory thermal optimum and field applicable enzyme is the central technical challenge that must be solved before the social cost analysis presented here can be applied to the full global footprint of biodegradable mulch film usage.

Despite this limitation, our results establish that single enzyme depolymerization of a chemically heterogeneous commercial polyester blend is achievable, identify the structural feature of the PETase fold that enables this, and quantify the social value that complete depolymerization captures relative to the partial fragmentation that current biodegradable mulch films deliver in soil. These findings reframe the design problem for chemically heterogeneous biodegradable plastics from one of enzyme combination to one of identifying enzymes whose cleft compatibility matches the targeted polymer, and provide a structural benchmark against which future ambient active polyesterases can be evaluated.

Rather than a simple transition toward biodegradable materials, blend approaches are required to manage the agricultural sustainable plastic usage. In particular, it recommends the substitution of non-biodegradable polymers to biodegradable polymers adapted to specific use conditions, while simultaneously establishing extended producer responsibility (EPR) schemes and strengthening recovery, recycling, and composting systems^6^. EPR or similar policy attempts would present opportunities for Kubu or similar approaches to reach minimum pollution from mulch film usages.

## Author Contribution

H.R. Kim - Conceptualization, Formal analysis, Methodology, Investigation, Visualization, Writing – original draft; H. Kim - Formal analysis, Methodology, Visualization, Writing - original draft; S. Jeong - Formal analysis, Visualization, Writing - original draft; J. H. Hwang - Investigation; D. Kim - Formal analysis; Y. W. Hong - Investigation; D.E. Suh - Funding acquisition; Sukkyoo Lee - Supervision, Writing - review and editing; Sangmin Lee - Supervision, Writing - review and editing; J.H. Cho - Methodology, Investigation, Validation, Writing - review and editing; J. Yu - Conceptualization, Formal analysis, Supervision, Visualization, Writing - original draft, Writing - review and editing; J. Oh - Conceptualization, Formal analysis, Supervision, Visualization, Funding acquisition, Writing - original draft, Writing - review and editing.

## Acknowledgements

We thank Central Instrument Facility of Kyung Hee University for their analytical assistance. This work was supported by the Ministry of SMEs and Startups (MSS, Republic of Korea) under the “Super Gap Startup 1000+ Project (20355745)”. This work was supported by National Research Foundation of Korea and Korea Basic Science Institute (RS-2024-00344054, RS-2025-25414953 and RS-2024-00401313),

## References

1 Lesk, C., Rowhani, P. & Ramankutty, N. Influence of extreme weather disasters on global crop production. Nature 529, 84–87, doi:10.1038/nature16467 (2016).

2 Tack, J., Barkley, A. & Nalley, L. L. Heterogeneous effects of warming and drought on selected wheat variety yields. Climatic Change 125, 489–500, doi:10.1007/s10584-014-1185-1 (2014).

3 Perry, E. D., Yu, J. & Tack, J. Using insurance data to quantify the multidimensional impacts of warming temperatures on yield risk. Nat Commun 11, 4542, doi:10.1038/s41467-020-17707-2 (2020).

4 Branca, G., Lipper, L., McCarthy, N. & Jolejole, M. C. Food security, climate change, and sustainable land management. A review. Agron Sustain Dev 33, 635–650, doi:10.1007/s13593-013-0133-1 (2013).

5 Zhang, L. et al. Plastic film mulching increases crop yields and reduces global warming potential under future climate change. Agr Forest Meteorol 349, doi:10.1016/j.agrformet.2024.109963 (2024).

6 FAO. Assessment of agricultural plastics and their sustainability: A call for action. In Assessment of Agricultural Plastics and Their Sustainability: A Call for Action. Rome. (2021).

7 Perego, G., Cella, G. D. & Bastioli, C. Effect of molecular weight and crystallinity on poly(lactic acid) mechanical properties. J Appl Polym Sci 59, 37–43, doi:Doi 10.1002/(Sici)1097-4628(19960103)59:1<37::Aid-App6>3.3.Co;2-7 (1996).

8 Burford, T., Rieg, W. & Madbouly, S. Biodegradable poly(butylene adipate-co-terephthalate) (PBAT). Phys Sci Rev 8, 1127–1156, doi:10.1515/psr-2020-0078 (2023).

9 Inkinen, S., Hakkarainen, M., Albertsson, A. C. & Sodergard, A. From lactic acid to poly(lactic acid) (PLA): characterization and analysis of PLA and its precursors. Biomacromolecules 12, 523–532, doi:10.1021/bm101302t (2011).

10 Wang, X., Peng, S. X., Chen, H., Yu, X. L. & Zhao, X. P. Mechanical properties, rheological behaviors, and phase morphologies of high-toughness PLA/PBAT blends by in-situ reactive compatibilization. Compos Part B-Eng 173, doi:10.1016/j.compositesb.2019.107028 (2019).

11 Sintim, H. Y. et al. In situ degradation of biodegradable plastic mulch films in compost and agricultural soils. Sci Total Environ 727, 138668, doi:10.1016/j.scitotenv.2020.138668 (2020).

12 Zhang, Y., Gao, W., Mo, A., Jiang, J. & He, D. Degradation of polylactic acid/polybutylene adipate films in different ratios and the response of bacterial community in soil environments. Environ Pollut 313, 120167, doi:10.1016/j.envpol.2022.120167 (2022).

13 Griffin-LaHue, D. et al. In-field degradation of soil-biodegradable plastic mulch films in a Mediterranean climate. Sci Total Environ 806, 150238, doi:10.1016/j.scitotenv.2021.150238 (2022).

14 Hao, Y. et al. Possible hazards from biodegradation of soil plastic mulch: Increases in microplastics and CO(2) emissions. J Hazard Mater 480, 136178, doi:10.1016/j.jhazmat.2024.136178 (2024).

15 Meyer-Cifuentes, I. E. et al. Synergistic biodegradation of aromatic-aliphatic copolyester plastic by a marine microbial consortium. Nat Commun 11, 5790, doi:10.1038/s41467-020-19583-2 (2020).

16 Yoshida, S. et al. A bacterium that degrades and assimilates poly(ethylene terephthalate). Science 351, 1196–1199, doi:10.1126/science.aad6359 (2016).

17 Tournier, V. et al. An engineered PET depolymerase to break down and recycle plastic bottles. Nature 580, 216–219, doi:10.1038/s41586-020-2149-4 (2020).

18 Yang, Y. et al. Complete bio-degradation of poly(butylene adipate-co-terephthalate) via engineered cutinases. Nat Commun 14, 1645, doi:10.1038/s41467-023-37374-3 (2023).

19 Williams, D. F. Enzymic hydrolysis of polylactic acid. Engineering in Medicine 10, 5–7 (1981).

20 Hajighasemi, M. et al. Biochemical and Structural Insights into Enzymatic Depolymerization of Polylactic Acid and Other Polyesters by Microbial Carboxylesterases. Biomacromolecules 17, 2027–2039, doi:10.1021/acs.biomac.6b00223 (2016).

21 Shalem, A., Yehezkeli, O. & Fishman, A. Enzymatic degradation of polylactic acid (PLA). Appl Microbiol Biotechnol 108, 413, doi:10.1007/s00253-024-13212-4 (2024).

22 Seo, H. et al. Landscape profiling of PET depolymerases using a natural sequence cluster framework. Science 387, eadp5637, doi:10.1126/science.adp5637 (2025).

23 Austin, H. P. et al. Characterization and engineering of a plastic-degrading aromatic polyesterase. Proc Natl Acad Sci U S A 115, E4350–E4357, doi:10.1073/pnas.1718804115 (2018).

24 Biundo, A. et al. Characterization of a poly(butylene adipate-co-terephthalate)-hydrolyzing lipase from Pelosinus fermentans. Appl Microbiol Biotechnol 100, 1753–1764, doi:10.1007/s00253-015-7031-1 (2016).

25 Avilan, L. et al. Concentration-Dependent Inhibition of Mesophilic PETases on Poly(ethylene terephthalate) Can Be Eliminated by Enzyme Engineering. ChemSusChem 16, e202202277, doi:10.1002/cssc.202202277 (2023).

26 Boyacı, İ. H. A new approach for determination of enzyme kinetic constants using response surface methodology. Biochemical Engineering Journal 25, 55–62 (2005).

27 Ghaffari-Moghaddam, M., Yekke-Ghasemi, Z., Khajeh, M., Rakhshanipour, M. & Yasin, Y. Application of response surface methodology in enzymatic synthesis: a review. Bioorg Khim 40, 275–285 (2014).

28 Hua, Z. et al. Quantitative analysis of PBAT microplastics and their degradation products in soil by mass spectrometry. Eco Environ Health 4, 100166, doi:10.1016/j.eehl.2025.100166 (2025).

29 Pan, H. et al. Isolation, characteristics, and poly(butylene adipate-co-terephthalate) (PBAT) degradation mechanism of a marine bacteria Roseibium aggregatum ZY-1. Mar Pollut Bull 201, 116261, doi:10.1016/j.marpolbul.2024.116261 (2024).

30 Guo, Z., Li, Y., Wang, M. & Ma, D. Catalytic Upcycling of PET: From Waste to Chemicals and Degradable Polymers. Acc Chem Res 58, 3184–3194, doi:10.1021/acs.accounts.5c00493 (2025).

31 Wu, W. M. & Criddle, C. S. Characterization of biodegradation of plastics in insect larvae. Methods Enzymol 648, 95–120, doi:10.1016/bs.mie.2020.12.029 (2021).

32 Kim, H. R. et al. Biodegradation of Polystyrene by Pseudomonas sp. Isolated from the Gut of Superworms (Larvae of Zophobas atratus). Environ Sci Technol 54, 6987–6996, doi:10.1021/acs.est.0c01495 (2020).

33 Corso, G., Stärk, H., Jing, B., Barzilay, R. & Jaakkola, T. Diffdock: Diffusion steps, twists, and turns for molecular docking. arXiv preprint arXiv:2210.01776 (2022).

34 Joo, S. et al. Structural insight into molecular mechanism of poly(ethylene terephthalate) degradation. Nat Commun 9, 382, doi:10.1038/s41467-018-02881-1 (2018).

35 Roth, C. et al. Structural and functional studies on a thermostable polyethylene terephthalate degrading hydrolase from Thermobifida fusca. Appl Microbiol Biotechnol 98, 7815–7823, doi:10.1007/s00253-014-5672-0 (2014).

36 Xiong, L., Jing, B., Chen, M. Y., Zheng, X. C. & Wu, W. Comprehensive environmental impact assessment of plastic film mulching with emphasis on waste disposal of discarded plastic film in sunflower production. J Clean Prod 404, doi:10.1016/j.jclepro.2023.136979 (2023).

37 Benavides, P. T., Lee, U. & Zarè-Mehrjerdi, O. Life cycle greenhouse gas emissions and energy use of polylactic acid, bioderived polyethylene, and fossil-derived polyethylene. J Clean Prod 277, doi:10.1016/j.jclepro.2020.124010 (2020).

38 EPA. EPA Report on the Social Cost of Greenhouse Gases: Estimates Incorporating Recent Scientific Advances. (2023).

39 Graf, M. et al. Differential effects of field-aged versus new LDPE and PLA/PBAT plastic film fragments on soil quality and crop productivity. J Hazard Mater 496, 139398, doi:10.1016/j.jhazmat.2025.139398 (2025).

40 Watkins, J. M., A., & Charles, D.. Annex1: The social cost of plastic-related harms. Minderoo Foundation: Nedlands, Australia. (2022).

41 Thompson, R. C. et al. Twenty years of microplastic pollution research-what have we learned? Science 386, eadl2746, doi:10.1126/science.adl2746 (2024).

42 Beaumont, N. J. et al. Global ecological, social and economic impacts of marine plastic. Mar Pollut Bull 142, 189–195, doi:10.1016/j.marpolbul.2019.03.022 (2019).

43 Fecker, T. et al. Active Site Flexibility as a Hallmark for Efficient PET Degradation by I. sakaiensis PETase. Biophys J 114, 1302–1312, doi:10.1016/j.bpj.2018.02.005 (2018).

44 Perz, V. et al. Substrate specificities of cutinases on aliphatic-aromatic polyesters and on their model substrates. N Biotechnol 33, 295–304, doi:10.1016/j.nbt.2015.11.004 (2016).

45 Gamerith, C. et al. Enzymatic Degradation of Aromatic and Aliphatic Polyesters by P. pastoris Expressed Cutinase 1 from Thermobifida cellulosilytica. Front Microbiol 8, 938, doi:10.3389/fmicb.2017.00938 (2017).

46 Ferrario, V., Pellis, A., Cespugli, M., Guebitz, G. & Gardossi, L. Nature Inspired Solutions for Polymers: Will Cutinase Enzymes Make Polyesters and Polyamides Greener? Catalysts 6, doi:10.3390/catal6120205 (2016).

47 Ribitsch, D. et al. Small cause, large effect: Structural characterization of cutinases from Thermobifida cellulosilytica. Biotechnol Bioeng 114, 2481–2488, doi:10.1002/bit.26372 (2017).

48 Knott, B. C. et al. Characterization and engineering of a two-enzyme system for plastics depolymerization. Proc Natl Acad Sci U S A 117, 25476–25485, doi:10.1073/pnas.2006753117 (2020).

49 Andersen, M., Kari, J., Borch, K. & Westh, P. Michaelis-Menten equation for degradation of insoluble substrate. Math Biosci 296, 93–97, doi:10.1016/j.mbs.2017.11.011 (2018).

50 Arnling Baath, J., Jensen, K., Borch, K., Westh, P. & Kari, J. Sabatier Principle for Rationalizing Enzymatic Hydrolysis of a Synthetic Polyester. Jacs Au 2, 1223–1231, doi:10.1021/jacsau.2c00204 (2022).

51 Kari, J. et al. A steady-state approach for inhibition of heterogeneous enzyme reactions. Biochem J 477, 1971–1982, doi:10.1042/BCJ20200083 (2020).

52 Madrid, B. et al. End-of-Life Management Options for Agricultural Mulch Films in the United States-A Review. Front Sustain Food S 6, doi:10.3389/fsufs.2022.921496 (2022).

53 Dong, H. T. et al. Recycling, disposal, or biodegradable-alternative of polyethylene plastic film for agricultural mulching? A life cycle analysis of their environmental impacts. J Clean Prod 380, doi:10.1016/j.jclepro.2022.134950 (2022).

