## Supplemental Tables and Figures for "PETase Kubu enables near-complete enzymatic depolymerization of commercial PLA/PBAT blend mulch film"

This PDF file includes:

Materials and Methods

Table S1-S9

Figures S1-S7

### Materials and Methods

#### Cloning and Purification of Plastic-Degrading Enzymes

##### Cloning of Plastic-Degrading Genes

Two types of genes, wild-type Kubu (genome accession number: QUNO01000019, complement: BCF44_11928) and wild-type PFL1 (genome accession number: AKVN01000044.1, complement: 237910 to 239118), were amplified by PCR using synthesized DNA sequences as templates. Each gene was amplified under gene-specific PCR conditions using appropriate primer pairs (Table S1). The PCR amplicons were digested with the restriction enzymes NdeI and XhoI and subsequently ligated into the pET-28b(+) expression vector. The resulting recombinant plasmids were transformed into *Escherichia coli* DH5α for plasmid propagation. Plasmids were extracted and verified by sequencing to confirm correct insertion and sequence integrity of the target genes. The verified expression constructs were then transformed into *E. coli* BL21(DE3) cells for protein expression and subsequent purification. Recombinant *E. coli* BL21(DE3) cells carrying the expression plasmids were cultured in 5 L of LB broth supplemented with kanamycin (15 mg/L) at 37 °C. Protein expression was induced at an OD600 of approximately 0.5 by adding isopropyl β-D-1-thiogalactopyranoside (IPTG) to a final concentration of 0.5 mM, followed by incubation at 18 °C for 48 h. Cells were harvested by centrifugation (4 °C, 4,000 rpm) for 20 min, and the resulting pellets were resuspended in lysis buffer. Cell disruption was performed by ultrasonication using repeated cycles of 1 s pulse and 2 s pause for a total processing time of 1 h. The lysates were subsequently centrifuged (13,000 rpm, 4 °C) for 1 h to remove insoluble debris, and the supernatants containing soluble recombinant proteins were collected for further purification.

##### Purification of Plastic-Degrading Enzymes

Recombinant target proteins were expressed as soluble forms and purified using immobilized metal affinity chromatography (IMAC). Because the target proteins contained multiple histidine tags, purification was performed using Ni–NTA agarose resin (Qiagen, Hilden, Germany), which exhibits high affinity toward poly-histidine residues. Clarified cell lysates were loaded onto a Ni–NTA column packed in an Econo-Column® chromatography column (Bio-Rad Laboratories Inc., CA, USA), pre-equilibrated with binding buffer. After washing to remove non-specifically bound proteins, the target proteins were eluted using a stepwise imidazole gradient ranging from 35 to 500 mM. Eluted fractions corresponding to imidazole concentrations of 7% to 100% were collected and analyzed.

To further improve protein purity, size-exclusion chromatography (SEC) was performed as a second purification step. The IMAC-purified protein fractions were concentrated and loaded onto a gel filtration chromatography (GFC) column connected to an ÄKTA go chromatography system (Cytiva, MA, USA) equilibrated with Tris-HCl buffer (50mM, pH8.0). Both proteins were purified using the same purification procedure. The purity and molecular weight of the recovered proteins were confirmed by SDS-PAGE analysis. Protein concentrations were determined using the Bradford assay.

#### Analysis of Enzymatic Plastic Degradation

##### Determination of Optimal Enzymatic Reaction Condition

Two enzymes with potential activity toward PLA/PBAT blend plastics were evaluated. To determine the optimal reaction temperature for each enzyme, degradation assays were conducted in 1 mL of Tris-HCl buffer (50mM, pH 8.0) containing PLA/PBAT (50:50, w/w) films at a concentration of 2.5 mg/mL and either Kubu or PFL1 at a final concentration of 0.1 mg/mL. Reactions were performed at 30, 45, 60, and 75 °C. The degradation efficiency was quantified by measuring the mass loss of the films after incubation. After enzymatic treatment, the plastic residues were collected by centrifugation at 8,000 rpm for 10 minutes. The supernatant was discarded, and the resulting pellet was recovered. The samples were then dried at room temperature for 24 h until no further change in weight was observed. The dried samples were weighed to determine weight loss, which was calculated based on the initial and final weights. All experiments were performed in triplicate.

After identifying the optimal temperature conditions, the effects of enzyme concentration on degradation efficiency were investigated. Kubu was tested at concentrations of 0.05, 0.10, 0.20, and 0.40 mg/mL, corresponding to the maximum concentration achievable after purification (0.40 mg/mL). PFL1 was evaluated at concentrations of 0.10, 0.20, 0.40, and 0.80 mg/mL, with 0.80 mg/mL representing the maximum obtainable concentration following purification. Film mass loss was measured to assess enzymatic degradation under each condition.

**Analysis of Apparent Enzymatic Degradation Kinetics**

To investigate whether degradation efficiency could be further enhanced through enzyme combination, assays were conducted using a mixture of the two enzymes that demonstrated effective PLA/PBAT degradation. PLA/PBAT blend films were incubated at concentrations of 2.5, 5.0, and 10.0 mg/mL in Tris-HCl buffer (50mM, pH 8.0) containing Kubu (0.1 mg/mL) and PFL1 (0.4 mg/mL). Reactions were carried out at 60 °C, and degradation efficiency was evaluated by measuring the mass loss of the films over time.

Apparent enzyme kinetics were evaluated by measuring the degradation rate of plastic substrates at varying substrate concentrations under a fixed enzyme concentration. The enzyme concentration was maintained at 0.1 mg/mL for all kinetic assays. Plastic substrates were added at final concentrations of 2.5, 5.0, 10.0, 20.0, and 40.0 mg/mL in the reaction mixtures. The reaction rates of plastic degradation were quantified by measuring weight loss within 5 hours. Apparent kinetic parameters were determined by fitting the initial rate data to a single rectangular hyperbolic Michaelis–Menten model using non-linear regression.

**Response Surface Methodology**

Response surface methodology (RSM) was employed to identify the precise optimal conditions for enzymatic degradation. RSM enables efficient experimental design and optimization in systems influenced by multiple variables, providing statistically robust response models that are more effective than conventional one-factor-at-a-time (OFAT) approaches[1,2]. Based on preliminary experiments, the optimization variables and their respective ranges were defined as follows: temperature (52.5-67.5 °C), pH (7.0-9.0), and agitation speed (0-200 rpm). The experimental design was constructed using a central composite design (CCD), and the center point was replicated three times to evaluate experimental reproducibility.

For each designed condition, Kubu was added at a final concentration of 10 μg/mL to a reaction mixture containing PLA/PBAT blend plastic (2.5 mg/mL) in 1 mL of Tris buffer adjusted to the designated pH. Reactions were conducted in a shaking incubator with controlled temperature and agitation speed. After 5 h of incubation, samples were collected, washed thoroughly with distilled water, dried, and weighed to determine the degradation rate based on mass loss. Statistical analysis of the experimental design and results was performed using JMP software (version 18) to construct a predictive response model. To validate the model, additional degradation experiments were conducted under the predicted optimal conditions using the same experimental procedures.

#### Characterization of Degraded Plastics

**Analysis of Degradation Products by LC-MS/MS**

To evaluate the characteristics of degraded plastics, enzymatic reactions with Kubu were performed under the optimal conditions determined by response surface methodology (RSM). The reactions were carried out in Tris-HCl buffer (50mM, pH 7.4) at 60 °C with agitation at 200 rpm. To analyze the degradation products generated by enzymatic reactions, liquid chromatography–tandem mass spectrometry (LC-MS/MS) was performed. Reaction supernatants obtained from two experimental groups; Kubu-treated and PFL1-treated were collected and filtered through 0.45 μm membrane filters prior to analysis. Chromatographic separation was carried out using an LC system equipped with a C18 reverse-phase column. The mobile phase consisted of water and acetonitrile (ACN), and a gradient elution program was applied, increasing the ACN content from 5% to 95%. Ionization was performed using electrospray ionization (ESI) in negative mode, and degradation products were detected and analyzed using a triple quadrupole mass spectrometer. Because lactic acid, the monomer of PLA, is difficult to reliably identify by LC-MS/MS under these conditions, its concentrations were quantified by high-performance liquid chromatography with diode array detection (HPLC-DAD) using an external standard calibration method. Standard solutions of lactic acid were analyzed to establish a calibration curve, and sample concentrations were determined by comparing their peak areas with those of the standards. HPLC-DAD analyses were performed at the Korea Polymer Testing & Research Institute.

**Gel Permeation Chromatography (GPC)**

The molecular weights of Kubu-treated plastics were analyzed by gel permeation chromatography (GPC) using an EcoSEC HLC-

8420 GPC system (Tosoh Bioscience, Tokyo, Japan). Samples were collected after 4 and 7 h of enzymatic reaction for analysis. The recovered samples were dissolved in chloroform ($CHCl_3$) to prepare solutions at a concentration of 3 mg/mL and filtered through 0.45 μm PTFE membrane filters to remove insoluble impurities. The mobile phase flow rate was set to 0.35 mL/min, and 10 μL of each sample solution was injected for analysis at 40 °C. Calibration curves for molecular weight determination were generated using polystyrene (PS) standards. GPC analyses were performed at the Korea Polymer Testing & Research Institute.

**Scanning Electron Microscopy (SEM)**

Surface morphological changes of the plastic films were examined using field-emission scanning electron microscopy (FE-SEM; Merlin Compact, Zeiss, Germany)[3]. The samples were mounted on SEM stubs using carbon tape and coated with a thin platinum layer prior to imaging. SEM images were acquired to compare untreated control samples with enzyme-treated samples after 4 and 7 h of incubation. However, samples treated for 7 h were highly fragmented, which prevented reliable surface morphology analysis.

#### Molecular Docking Simulation

**Enzyme Structures Preparation**

Crystal structures of Kubu PETase (PDB: 8YTW), wild-type IsPETase (PDB: 5XJH), and wild-type TfCut (PDB: 4CG1) were used for docking. Chain A was retained; water molecules and non-protein heteroatoms were removed. No energy minimization or side-chain repacking was applied. Co-crystal structures of IsPETase-MHET (PDB: 7XTW) and TfCut-MHET (PDB: 7XTV) were used as references for active site geometry comparison. As these co-crystal structures were solved using catalytic serine mutants (S160A and S170A, respectively), docking was performed against the corresponding wild-type structures.

**Ligand Library Preparation**

Ligand structures were prepared from canonical SMILES using RDKit[4]. The library included L-form PLA oligomers (2~4-mer) with hydroxyl and carboxyl termini; PBAT model oligomers (BAB, BTB, and BABTB) constructed with 1,4-butanediol-capped termini, based on Yang et al.[5]; and BHET as a PET-derived substrate (Table S4; Figure S6). BHET was included to assess whether Kubu accommodates PET-derived substrates in a catalytically relevant pose, and to allow comparison with IsPETase and TfCut under identical docking conditions.

**Docking and Pose Filtering**

Molecular docking was performed using DiffDock v1.1[6] on an NVIDIA RTX 4080 GPU, generating 1,000 poses per enzyme-ligand pair under the default blind-docking protocol. All poses were subsequently filtered through an identical four-stage pipeline (Table S6), with enzyme-specific parameters limited to oxyanion-hole residue identities and catalytic serine position (Table S5). Each pose was prepared as a protein–ligand complex PDB file using PyMOL[7] and analyzed with PLIP[8] to identify non-covalent interactions. Retention required that both oxyanion-hole NH residues simultaneously donate a backbone hydrogen bond to a ligand oxygen; poses satisfying only one such contact were discarded. Surviving poses were then assessed for physical plausibility, with protein–ligand steric clashes, distorted ligand geometries, and anomalous bond lengths each serving as independent rejection criteria. Remaining poses were evaluated for catalytic geometry, requiring an Oγ–ester carbonyl C distance of 1.5–4.0 Å and an Oγ–C=O angle of 70–120°. Ester carbonyl carbons were identified using the SMARTS pattern [CX3](=[OX1])-[OX2]-[#6],where multiple valid candidates existed, the carbon with the smallest combined deviation from the midpoint of each range was selected. Final poses were visually inspected in PyMOL to confirm productive orientation of the leaving-group oxygen toward the catalytic histidine.

#### Estimating Social Cost of Alternative Mulching Strategies

**Social cost of PE mulching film: GHG emission during incineration or landfill**

The social cost of PE mulch film disposal is estimated using weighted average emission factors from landfill and incineration pathways reported by Xiong et al. (2023)[9] and Benavides et al. (2020)[10]. The weighted averages reflect disposal pathway shares reported by OECD (2022)[11], FAO (2021)[12], and Xiong et al. (2023)[9]. Carbon price scenarios are based on EPA (2023)[13].

$$SC_{PE} = \left( \sum_{i=landfill\ or\ incineration} Disposal\ ratio_i \times Emission\ factor_i \right) \times Carbon\ price$$

**Social cost of BDM mulching film: microplastic residues from incomplete degradation**

Residual microplastic emissions from PLA and PBAT mulch films are based on Hao et al. (2024)[14]. Hao et al. (2024) reported the amount of microplastic emissions generated when all mulching film used in China are replaced with PLA or PBAT, respectively. The social cost of microplastic pollution includes both human health damages and ecosystem service losses. Health related damages are estimated using the Disability-Adjusted Life Year and its value (Watkins et al. 2022)[15]. The unit social cost of health is calculated by dividing the estimated annual health-related social cost of micro/nano-plastic exposure ($28.5 billion) by the estimated annual emissions of microplastics to the environment (10 to 40 million tons)[16]. Due to the limited availability of empirical estimates on agricultural ecosystem damages associated with microplastic pollution, the estimated value of marine ecosystem loss is used as a proxy for ecosystem related damages[17].

$$SC_{BDM} = Emission\ factor_{j:\ PLA\ or\ PBAT} \times (unit\ of\ SC\ of\ health + Ecosystem\ loss)$$

**Table S1. Enzyme kinetics analysis of Kubu**

| | $K_{m\_app}$ (mg·mL⁻¹) | $V_{max\_app}$ (mg·mL⁻¹·5 h⁻¹) |
|---|---|---|
| **PETase** | 9.37 | 11.49 |

**Table S2. Experimental design for response surface methodology**

| Design | Temperature | pH | RPM |
|---|---|---|---|
| --- | 52.5 | 7.0 | 0 |
| --+ | 52.5 | 7.0 | 100 |
| -00 | 52.5 | 8.0 | 200 |
| -+- | 52.5 | 9.0 | 0 |
| -++ | 52.5 | 9.0 | 100 |
| 0-0 | 60 | 7.0 | 200 |
| 00- | 60 | 8.0 | 0 |
| 000 | 60 | 8.0 | 100 |
| 000 | 60 | 8.0 | 100 |
| 000 | 60 | 8.0 | 100 |
| 00+ | 60 | 8.0 | 200 |
| 0+0 | 60 | 9.0 | 100 |
| +-- | 67.5 | 7.0 | 0 |
| +-+ | 67.5 | 7.0 | 200 |
| +00 | 67.5 | 8.0 | 100 |
| ++- | 67.5 | 9.0 | 0 |
| +++ | 67.5 | 9.0 | 200 |

**Table S3. Quantitative analysis of plastic degradation products (lactic acid)**

| | Lactic acid (mg·L⁻¹) |
|---|---|
| PETase | 257 |
| Lipase | 261 |
| PETase+Lipase | 258 |

**Table S4. SMILES structures of the ligand library used in molecular docking**

| Ligand | SMILES |
|---|---|
| **PLA2** | OC(=O)[C@H](C)OC(=O)[C@H](C)O |
| **PLA3** | OC(=O)[C@H](C)OC(=O)[C@H](C)OC(=O)[C@H](C)O |
| **PLA4** | OC(=O)[C@H](C)OC(=O)[C@H](C)OC(=O)[C@H](C)OC(=O)[C@H](C)O |
| **PBAT(BAB)** | OCCCCOC(=O)CCCCC(=O)OCCCCO |
| **PBAT(BTB)** | OCCCCOC(=O)c1ccc(C(=O)OCCCCO)cc1 |
| **PBAT(BABTB)** | OCCCCOC(=O)CCCCC(=O)OCCCCOC(=O)c1ccc(C(=O)OCCCCO)cc1 |
| **BHET** | C1=CC(=CC=C1C(=O)OCCO)C(=O)OCCO |

**Table S5. Oxyanion-hole residues and catalytic serine positions used in pose filtering.**

| Enzyme | Oxyanion residue 1 | Oxyanion residue 2 | Catalytic serine |
|---|---|---|---|
| **Kubu PETase (8YTW)** | F92 (backbone NH) | M163 (backbone NH) | S162 |
| **IsPETase (5XJH)** | Y87 (backbone NH) | M161 (backbone NH) | S160 |
| **TfCut (4CG1)** | Y100 (backbone NH) | M171 (backbone NH) | S170 |

**Table S6. Pose filtering criteria.**

| Stage | Criterion |
|---|---|
| **1. Oxyanion hole** | Both oxyanion-hole NH residues must simultaneously H-bond to a ligand oxygen; poses with only one contact discarded |
| **2. Pose quality** | No protein–ligand heavy-atom pair < 1.50 Å; no non-bonded ligand pair < 1.20 Å; fewer than 2 pairs < 0.50 × ΣvdW radii; all bonds within 0.50–2.00 × Σcovalent radii; no inter-fragment distance < 1.20 Å |
| **3. Catalytic geometry** | Ester carbonyl by SMARTS [CX3](=[OX1])-[OX2]-[#6]; Ser Oγ–C distance 1.5–4.0 Å; Oγ–C=O angle 70–120° |
| **4. Visual inspection** | Ser Oγ toward ester carbonyl carbon; leaving-group O toward catalytic histidine; oxyanion-hole geometry confirmed |

**Table S7. Number of productive docking poses retained after filtering (1,000 poses per enzyme–ligand pair).**

| Ligand | Kubu (8YTW) | IsPETase (5XJH) | TfCut (4CG1) |
|---|---|---|---|
| **PLA2** | 77 | 381 | 29 |
| **PLA3** | 145 | 769 | 248 |
| **PLA4** | 34 | 590 | 197 |
| **PBAT BAB (adipate unit)** | 311 | 379 | 569 |
| **PBAT BTB (terephthalate unit)** | 311 | 455 | 227 |
| **PBAT BABTB** | 68 | 140 | 116 |
| **BHET** | 662 | 130 | 75 |

**B, 1,4-butanediol; A, adipate; T, terephthalate; BHET, bis(2-hydroxyethyl) terephthalate.**

**Table S8. Parameters used for estimating the social cost of PE mulch film disposal**

| Parameter | Value | Unit | Source |
|---|---|---|---|
| Disposal ratio | 59 (Landfill)<br>19 (Incineration) | % | OECD (2022), FAO (2021)<br>Xiong et al. (2023), |
| Emission factor | 0.7 (Landfill)<br><br>2.9 (Incineration) | tCO2eq/ton of PE | Xiong et al. (2023),<br><br>Benavides et al. (2020) |
| Carbon price | 120, 190, 340 | $/tCO2eq | EPA (2023) |

**Table S9. Parameters used for estimating the social cost of Biodegradable mulch film**

| Parameter | Value | Unit | Source |
|---|---|---|---|
| Emission factor | 0.02 (PLA)<br>0.11 (PBAT) | t/ton of mulching film | Hao et al. (2024) |
| Unit of Social cost of health | 712.5 ~ 2,850 | $/ton | Watkins et al. (2022)<br>Thompson et al. (2024) |
| Ecosystem loss | 3,300 ~ 33,000 | $/ton | Beaumont et al. (2019) |

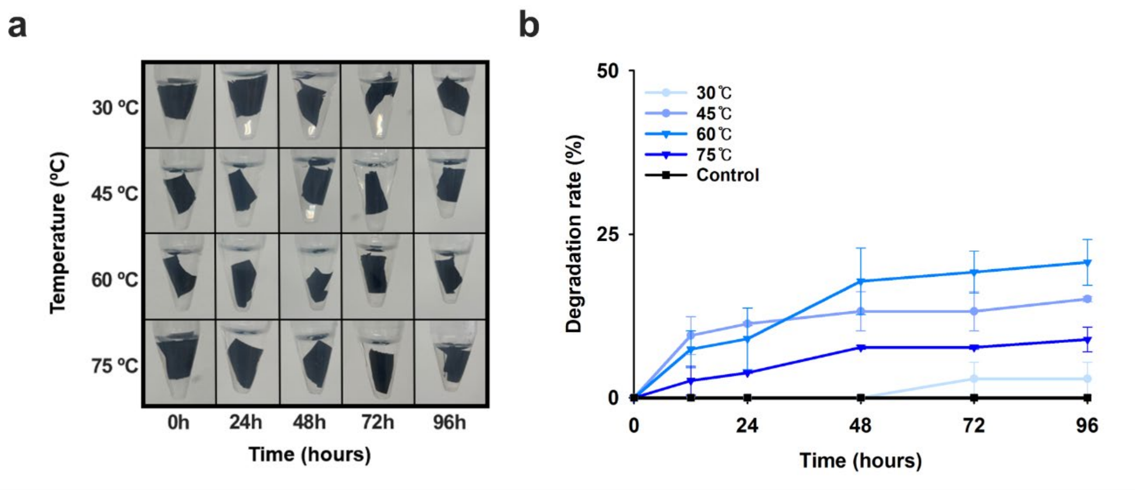


**Figure S1. Temperature-dependent degradation of PLA/PBAT blend films by PFL1.** (a) Time-course degradation of PLA/PBAT blend films by PFL1 at different temperatures. (b) Degradation efficiency of PLA/PBAT blend films as a function of temperature and reaction time for PFL1.

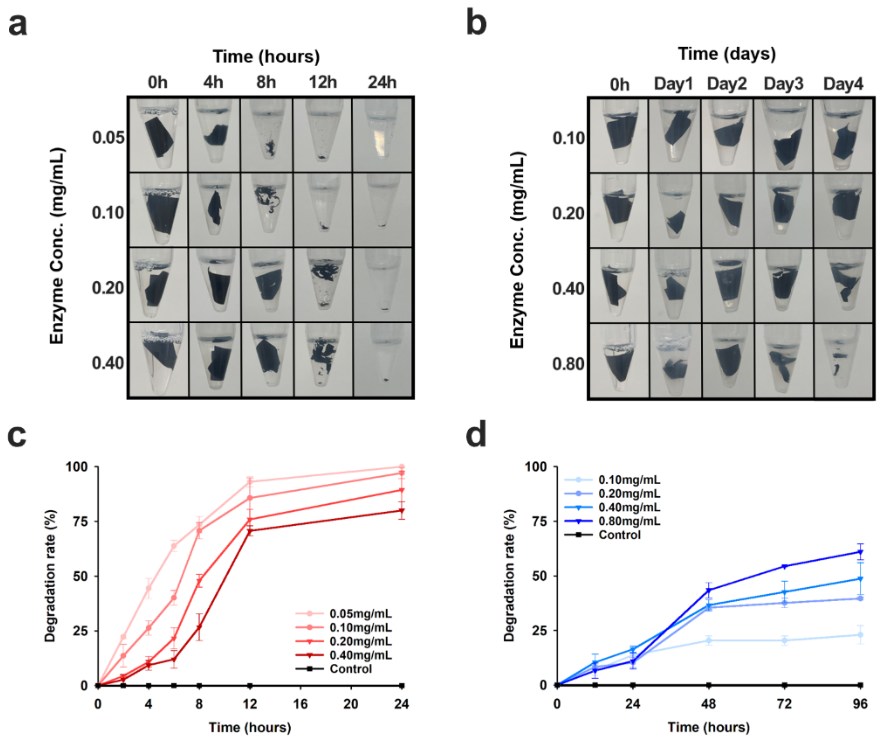


**Figure S2. Effect of enzyme concentration on degradation of PLA/PBAT blend films.** (a,b) Time-course degradation of PLA/PBAT blend films by Kubu (a) and PFL1 (b) at different enzyme concentrations. (c,d) Degradation efficiency of PLA/PBAT blend films as a function of enzyme concentration and reaction time for Kubu (c) and PFL1 (d).

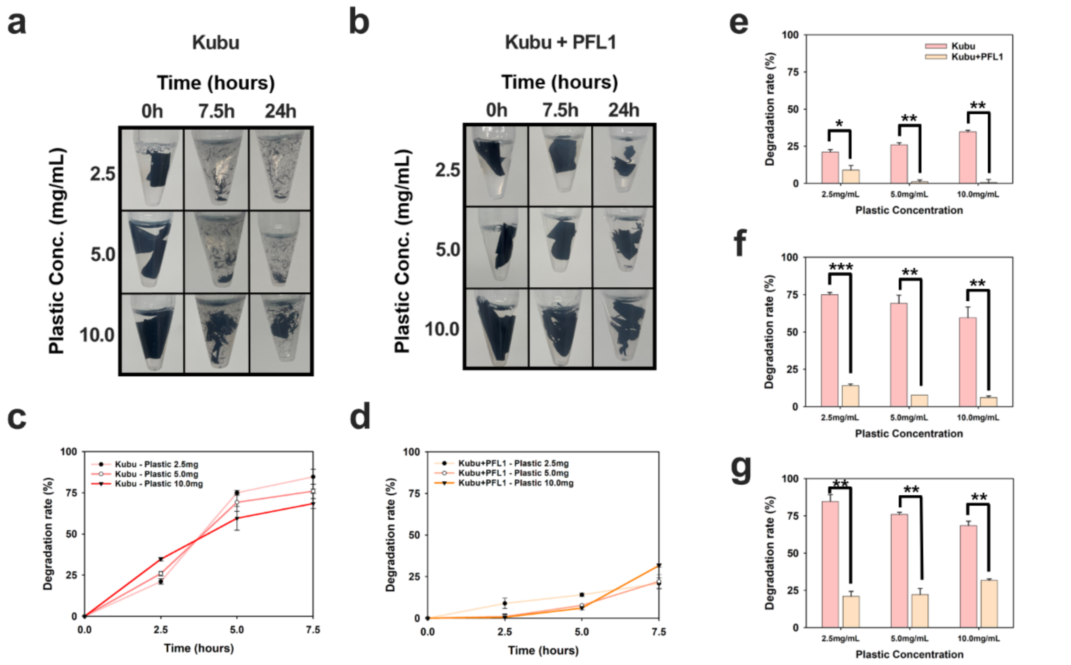


**Figure S3. Effect of enzyme combinations on degradation of PLA/PBAT blend films.** (a,b) Time-course degradation of PLA/PBAT blend films by Kubu (PETase) alone (a) and by a combination of Kubu (PETase) and PFL1 (Lipase) (b). (c,d) Degradation efficiency of PLA/PBAT blend films by Kubu alone (c) and by the Kubu–PFL1 combination (d). (e–g) Comparison of degradation efficiency at 2.5 h (e), 5 h (f), and 7.5 h (g). Statistical significance is indicated as $p < 0.05$ (*), $p < 0.01$ (**), and $p < 0.001$ (***).

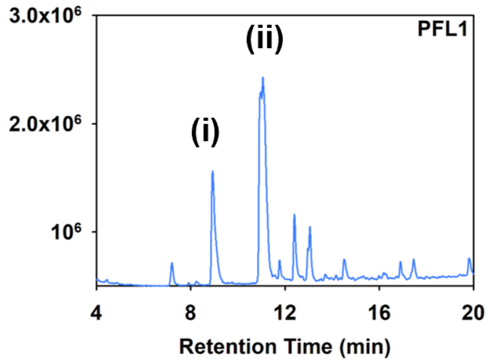


**Figure S4. LC–MS/MS chromatograms of degradation products.**

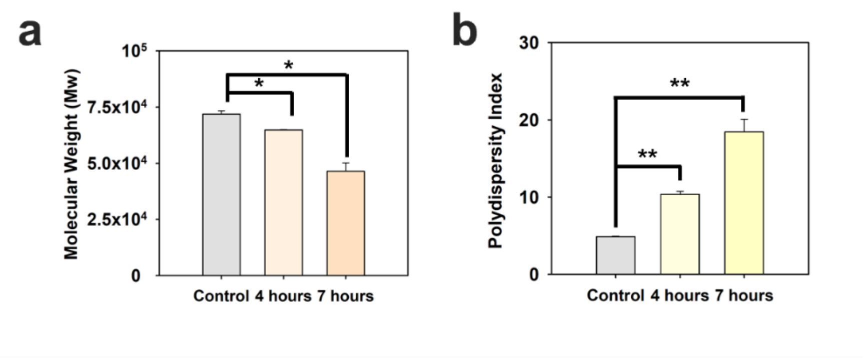


**Figure S5. Changes in weight-average molecular weight (a) and polymer dispersity index (b) during enzymatic degradation.**

PLA2

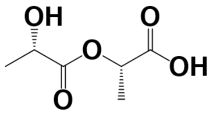

PLA3

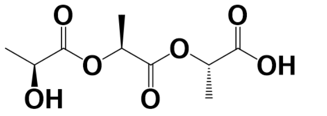

PLA4

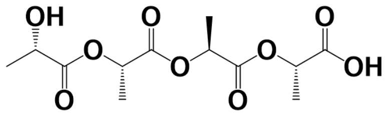

BHET

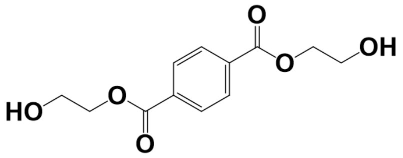

PBAT(BAB)

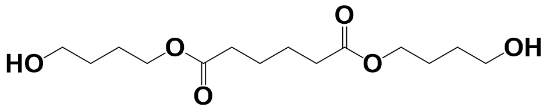

PBAT(BTB)

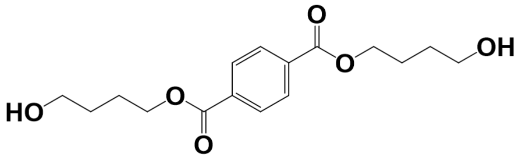

PBAT(BABTB)

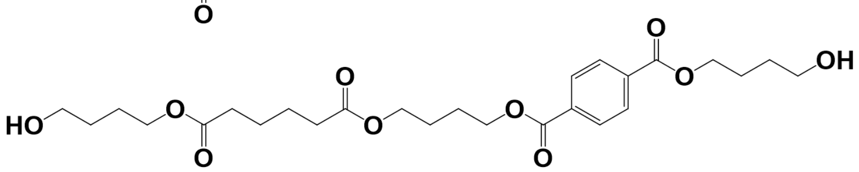

**Figure S6. 2D structures of ligands used in molecular docking.** L-form PLA oligomers (PLA2, PLA3, PLA4) with hydroxyl and carboxyl termini; BHET as a PET-derived model substrate; and PBAT model oligomers (BAB, BTB, BABTB) with 1,4-butanediol-capped termini.

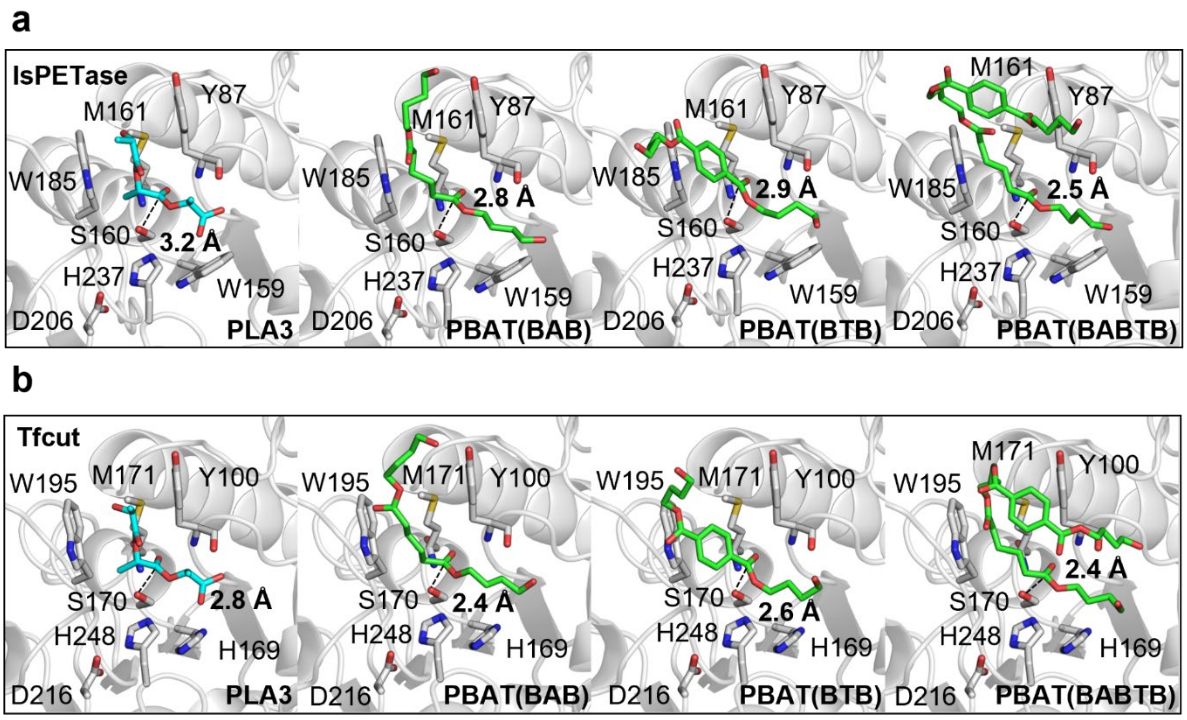


**Figure S7. Representative productive docking poses of PLA and PBAT ligands in the active sites of IsPETase (top) and TfCut (bottom).** The catalytic triads Ser160–His237–Asp206 (IsPETase) and Ser170–His248–Asp216 (TfCut) are shown. The backbone amides of Y87 and M161 (IsPETase) or Y100 and M171 (TfCut) form the oxyanion hole. PLA3 is shown in cyan; PBAT oligomers (BAB, BTB, BABTB) are shown in green. Dashed lines indicate the distance between the catalytic serine Oγ and the scissile ester carbonyl carbon.

**SI references**


1 Ghaffari-Moghaddam, M., Yekke-Ghasemi, Z., Khajeh, M., Rakhshanipour, M. & Yasin, Y. Application of response surface methodology in enzymatic synthesis: a review. *Bioorg Khim* **40**, 275–285 (2014).

2 Boyaci, I. H. A new approach for determination of enzyme kinetic constants using response surface methodology. *Biochem Eng J* **25**, 55–62 (2005). https://doi.org/10.1016/j.bej.2005.04.001

3 Kim, H. R. *et al.* Isolation of a polyethylene-degrading bacterium, Acinetobacter guillouiae, using a novel screening method based on a redox indicator. *Heliyon* **9**, e15731 (2023). https://doi.org/10.1016/j.heliyon.2023.e15731

4 RDKit: Open-source cheminformatics (Zenodo, 2026).

5 Yang, Y. *et al.* Complete bio-degradation of poly(butylene adipate-co-terephthalate) via engineered cutinases. *Nat Commun* **14**, 1645 (2023). https://doi.org/10.1038/s41467-023-37374-3

6 Corso, G., Stärk, H., Jing, B., Barzilay, R. & Jaakkola, T. in *International Conference on Learning Representations* (2023).

7 Schrödinger, LLC. *The PyMOL Molecular Graphics System, Version 3.1.0* (2025).

8 Schake, P., Bolz, S. N., Linnemann, K. & Schroeder, M. PLIP 2025: introducing protein-protein interactions to the protein-ligand interaction profiler. *Nucleic Acids Res* **53**, W463–W465 (2025). https://doi.org/10.1093/nar/gkaf361

9 Xiong, L., Jing, B., Chen, M. Y., Zheng, X. C. & Wu, W. Comprehensive environmental impact assessment of plastic film mulching with emphasis on waste disposal of discarded plastic film in sunflower production. *J Clean Prod* **404** (2023). https://doi.org/10.1016/j.jclepro.2023.136979

10 Benavides, P. T., Lee, U. & Zarè-Mehrjerdi, O. Life cycle greenhouse gas emissions and energy use of polylactic acid, bio-derived polyethylene, and fossil-derived polyethylene. *J Clean Prod* **277** (2020). https://doi.org/10.1016/j.jclepro.2020.124010

11 OECD. Global Plastics Outlook: Economic Drivers, Environmental Impacts and Policy Options. (Paris, 2022).

12 FAO. Assessment of agricultural plastics and their sustainability: A call for action. (Rome, 2021).

13 U.S. Environmental Protection Agency. Report on the Social Cost of Greenhouse Gases: Estimates Incorporating Recent Scientific Advances. (National Center for Environmental Economics; Office of Policy; Climate Change Division; Office of Air and Radiation, Washington, DC, 2023).

14 Hao, Y. *et al.* Possible hazards from biodegradation of soil plastic mulch: Increases in microplastics and CO(2) emissions. *J Hazard Mater* **480**, 136178 (2024). https://doi.org/10.1016/j.jhazmat.2024.136178

15 Watkins, J., Charles, D. & Merkl, A. Annex 1: The Social Cost of Plastic-Related Harms. (Minderoo Foundation, Nedlands, Australia, 2022).

16 Thompson, R. C. *et al.* Twenty years of microplastic pollution research-what have we learned? *Science* **386**, eadl2746 (2024). https://doi.org/10.1126/science.adl2746

17 Beaumont, N. J. *et al.* Global ecological, social and economic impacts of marine plastic. *Mar Pollut Bull* **142**, 189–195 (2019). https://doi.org/10.1016/j.marpolbul.2019.03.022
